# Visual exoproteomics of *Clostridium thermocellum* during anaerobic biomass-degradation identifies functional spirosomes

**DOI:** 10.64898/2026.05.21.726627

**Authors:** Matthew P. Agdanowski, Matthew J. Kensil, Trevor H. Moser, Ethan Humm, Yaneli I. Guandique, Kayleigh Mason-Chalmers, Tracy al-Set, Rachel R. Ogorzalek Loo, James E. Evans, Robert P. Gunsalus, Joseph A. Loo, Jose A. Rodriguez

## Abstract

Visual proteomics enables the study of low-abundance proteins and identification of unknown complexes from heterogeneous samples by complementing high-resolution cryogenic electron microscopy (cryoEM) with external inputs on protein identity such as mass spectrometry. Using this approach, we interrogated the exoproteome of the anaerobic cellulose-degrading bacterium *Clostridium thermocellum* as it carried out biomass degradation. Mass spectrometry indicated a broad exoproteome composition, including cellulose degrading machinery CelA and CipA. A focus on large exoproteome assemblies revealed abundant protein filaments and pleomorphic vesicular structures. Analysis of the most abundant protein filaments yielded an ∼4 □ resolution native structure that, aided by mass spectrometry, *de novo* modeling, and structural searching, was found to be the aldehyde-alcohol dehydrogenase (AdhE) spirosome. AdhE contained both NAD^+^ and Fe in their expected binding sites and biochemical and structural analyses of enriched spirosome preparations indicated they were functional. Altered NADH solution concentrations triggered conformational changes in the exoproteomic spirosomes, and the constituent AdhE remained capable of ethanol production. Although the basis for functional extracellular spirosome accumulation in live anaerobic *C. thermocellum* cultures remains unclear, their abundance in crude exoproteomes suggests their presence could influence biomass fueled *C. thermocellum* growth.

## Introduction

*Clostridium thermocellum* (*Ct*), an anaerobic gram-positive bacterium ubiquitous in hot, cellulose-rich environments, is a model organism for its high rate of biomass degradation^1^. Interest in *Ct* is driven by its ability to degrade cellulose, a highly crystalline polymer that resists enzymatic breakdown^2,3^, which can meet a need for renewable, cost-efficient energy and commodity production^4,5^. Improved understanding of the bioprocessing ability of *Ct* could therefore inform potential platforms for biomanufacturing^6^, particularly given the high abundance of cellulosic biomass, an untapped resource^3,7^.

*Ct* and other anaerobic bacteria and archaea achieve high metabolic rates in part by expressing a host of carbohydrate degrading enzymes, CAZymes^8^. Some of these enzymes organize into a multienzyme complex known as the cellulosome^9,10^, an extracellular assembly of a scaffolding protein that tethers the cellulolytic CAZymes by interaction of cohesin and dockerin domains^11,12^. Cellulosomes are among the most elaborate enzymatic systems found in bacteria^13^ and are commonly depicted as cell-surface-anchored complexes, but studies have shown that *Ct* can also produce cell-free cellulose-degrading components untethered to the bacterial cell wall that can diffuse away from the cell to degrade polysaccharide substrates^14–16^. To date, high resolution structural work on cellulosomes and their components has relied on X-ray crystallography or NMR^17–20^, with cryoEM only recently providing limited insights^21^. In addition to the cellulosome, *Ct* expresses a wide array of extracellular proteinaceous complexes, most involved in biomass sensing, gene regulation, and metabolism^22,23^. As the structures of these exoproteomic components remain understudied, a comprehensive understanding of the molecular workings of *Ct*-associated extracellular complexes is relevant for further improvement of biomass degradation.

By facilitating the structural characterization of unknown protein complexes directly from impure biological systems such as cell lysates and even intact cells, visual proteomics offers a powerful approach to interrogate exoproteomes^24–27^. This is especially true given its ability to identify unknown or unannotated protein densities observed in cryoEM data without the need for chemical fixation or labeling^28,29^. Its success has been achieved by a dramatic expansion in the capabilities of both cryoEM and mass spectrometry, driven in part by the development of direct electron detectors, improvements in microscope stability, and algorithmic advances^30–33^. Mass spectrometry-based proteomics is an excellent complement to cryoEM, enabling the high-throughput identification and quantification of proteins in complex biological samples, thereby providing a comprehensive inventory of the molecular components present in the sample^34,35^. Furthermore, mass spectrometry can also provide additional information about protein abundance, stoichiometry, post-translational modifications, and interaction partners, which can constrain candidate identities and guide model-building or structure prediction^36–38^.

Leveraging the strengths of visual proteomics^39^, we applied the approach to study high molecular weight extracellular protein complexes derived from *Ct* during its anaerobic growth on cellobiose-containing media (Figure S1). Aiming to further insights into cellulose degradation-associated complexes^40^, we interrogated high molecular weight fractions of the *Ct* exoproteome to reveal varied proteinaceous species including regularly twisting protein filaments. A combination of cryoEM, cryoET, mass spectrometry and computational modeling revealed the filaments to be spirosomes, composed of the aldehyde-alcohol dehydrogenase AdhE^41^. Biochemical analyses indicated the spirosomes were functional as isolated and that their component AdhE monomers retained NAD^+^ and iron cofactors. Their presence and function in the extracellular milieu of biomass-degrading *Ct* cultures suggests they may play a structurally regulated extracellular function^42,43^.

## Results

### Analysis of Ct exoproteome during biomass degradation

Hypothesizing that cellulose-degrading and related protein complexes are present in the extracellular media of *Ct* cultures during anaerobic growth on the disaccharide cellobiose, we harvested late log phase cultures (Figure S1) and isolated their high molecular weight exoproteomic fractions. Species with a molecular weight greater than 100 kDa were concentrated and subjected to gel filtration, showing evidence of proteinaceous assemblies (Figure 1A). Upon inspection by negative stain electron microscopy (nsEM), the fractions showed a variety of complexes including filaments, vesicles, irregular oligomers, small clusters of globular proteins, and the occasional flagellum (Figure 1B). Each fraction was further subjected to bottom-up proteomics to identify the protein components (Figure 1C). Discrete bands were evident by SDS-PAGE, indicating that the observed nsEM complexes corresponded to a small set of protein complexes (Figure S2). Within each analyzed fraction, all of the highly abundant proteins were associated with cellulose degradation, with the top protein hits corresponding to cellobiodases, xylanases, and endoglucanases (Table S1).

**Figure 1.**
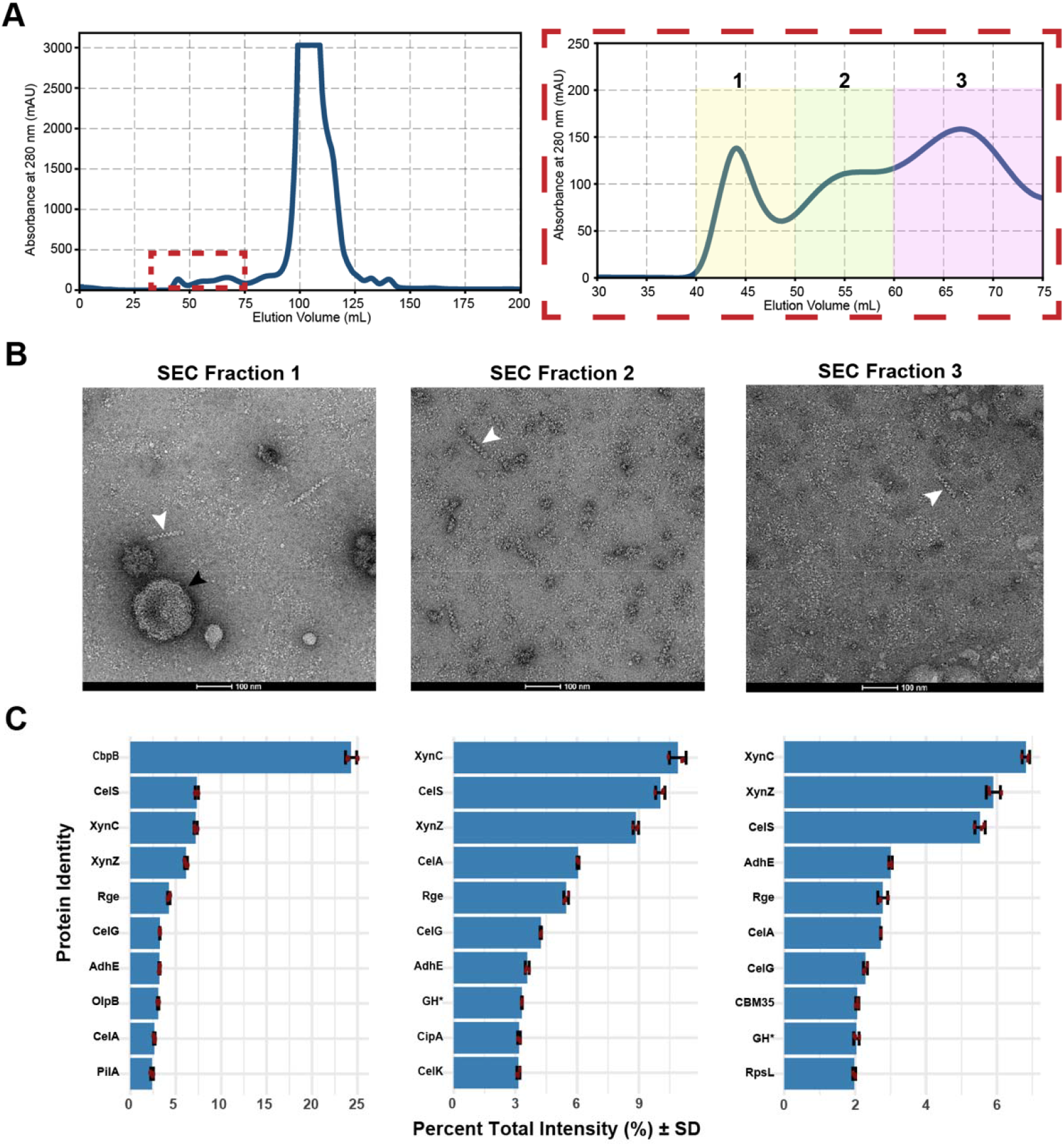
Investigation of *C. thermocellum* exoproteome. (A) Size fractionation of *Ct* growth media reveals large molecular weight species, with fractions indicating major species identified by peaks in the chromatogram (inset). (B) Several large species, filaments (white arrows) and vesicles (black arrows) are observed by nsEM. (C) Bottom-up proteomics for each sample shows proteins implicated in biomass degradation.

One fraction was found to predominantly contain isolated vesicles that could be further enriched by ultracentrifugation. In nsEM micrographs, the vesicles appeared pleomorphic and of various sizes (Figure S3A); by cryoEM they appeared to have membranes decorated with discrete protein complexes (Figure S3B). Tomograms of the vesicles further revealed a range of morphologies and shapes that deviated from an ideal spherical geometry (Figure S4A-I), with an average diameter of ∼28 nm and a mean cross-sectional area of ∼2700 nm^2^ (Figure S4J,K). Proteomic analysis of this fraction identified the most abundant protein as the sugar transport protein CbpB (Figure S4L, Table S2), which is responsible for cellodextrin uptake and is a critical component of the cellulosome^44–46^. Interestingly, CbpB was absent from fractions devoid of vesicles, suggesting that it may be vesicle-associated.

Also present in several high molecular weight exoproteome fractions were filamentous assemblies approximately 10 nm in diameter and ranging from 50 nm to over 200 nm long (Figure S5). These filaments were further enriched by centrifugation and subjected to proteomic analysis, revealing, among the most abundant protein, a cytosolic enzyme implicated in ethanol production: aldehyde-alcohol dehydrogenase (AdhE) (Table S3). Although its extracellular abundance was a surprise, the presence of AdhE^47^ agreed with its facilitation of ethanol production by *Ct*^*41*^. Despite the abundance and regularity of the filamentous assemblies, their identity was not clear from nsEM micrographs alone, prompting investigation by single particle cryoEM.

### Reconstruction of exoproteomic Ct filaments by single particle cryoEM

Enriched filament fractions were subjected to single particle cryoEM, with the expectation that their regular twist would enable helical reconstruction. Proteomic analysis of fractions devoid of large vesicles and smaller protein aggregates (Figure S6A), confirmed that they retained the same cellulolytic enzymes observed in the crude high molecular weight fractions (Figure 2A). Their imaging by cryoEM revealed a sufficiently high concentration of filaments for helical reconstruction (Figure S6B), which was attempted from an initial set of 4,863 high-resolution cryoEM micrographs. These were processed to yield 94,278 particles that were organized into distinct 2D classes whose features matched those observed in nsEM micrographs (Figure 2B,C); the best quality classes were selected for 3D *ab initio* reconstruction. Helical refinement from these particles resulted in a map at ∼4.07 □ (Figure 2D) of filaments with a twist of 193.97° and rise of 60.06 □. That map was suitable for structure identification and *de novo* model building (Figure S7, Table S4).

**Figure 2.**
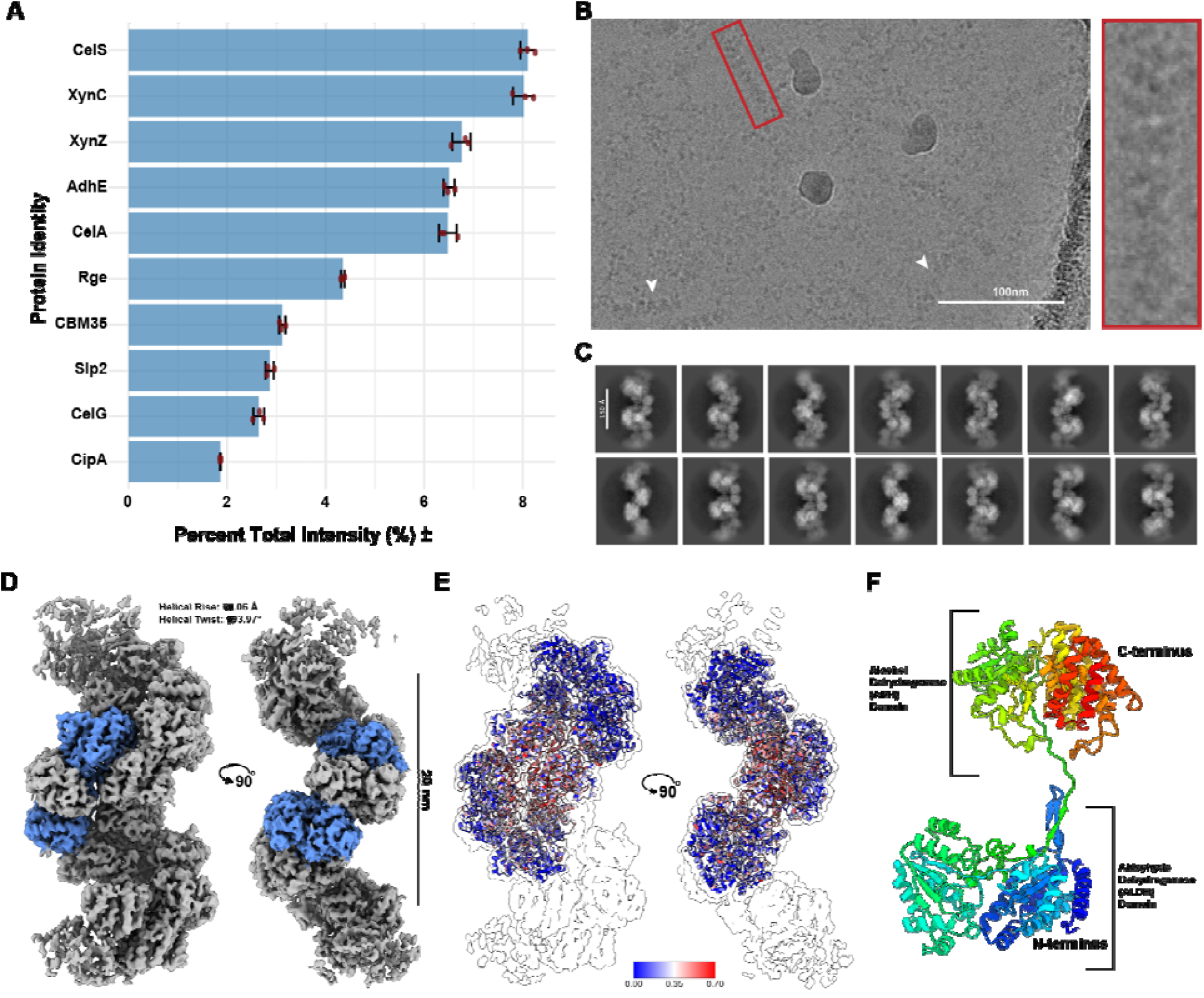
Identification of unknown filament identity. (A) Proteomics of enriched filaments reveals a protein list of potential filament identities ranked by total protein intensity. (B) CryoEM micrographs show well behaved filaments used in high-resolution data collection. (C) Resulting 2D class averages with discernible features. (D) Helical reconstruction results in a map of a twisting filament, with spirosome monomer highlighted in blue. (E) Q-score analysis of AdhE (PDB: 8UHW) structure confirms best fit of potential proteins. (F) Cartoon model of AdhE monomer displaying a two-domain architecture.

### Ct exoproteome filaments are composed of AdhE

A panel of 32 AlphaFold2^48^ models and published structures corresponding to the 10 most abundant *Ct* proteins in the enriched filament fraction was assessed for fit into the helical reconstruction map density. For each, quality of fit was determined by the fraction and total number of model atoms that matched the map (Figure S8A, Tables S5-8). This analysis indicated that the best matching model was of *Ct* AdhE, with 64.16%, or 19,755 of its atoms across six protein chains fitting the map density. Fitting the deposited AdhE structure (PDB: 8UHW) to the map yielded a mean Q-score of 0.25 across all six of its chains (Figure 2E). The same analysis performed on a single monomer of the AdhE protein resulted in a marginally better Q-score of 0.31 (Figure S9A). However, relaxing the pose for either the single monomer or entire six chain complex in Rosetta^49,50^ nearly doubled the model-to-map correlation, as reflected by improved Q-scores of 0.54 for the monomer and 0.51 for the entire complex (Figure S9B,C). The Q-scores for other top candidate proteins in the set, such as the small cellulosomal-scaffolding protein A (PDB: 1ANU), yielded lower mean Q-scores (Figure S8B), indicating that they were poorer matches.

In parallel, automated model-building was used to trace chains within the structure *ab initio* and supply a template for structure-based similarity searches that might identify the target protein. The unsharped map from helical processing was provided as an input to the ModelAngelo program in sequence-independent mode^51^. A resulting fragmented model was joined to produce a single chain fragment to be used in structural searches (Figure S10A,B). A structural comparison of this model against the DALI database^52^ indicated its high similarity to the *Ct* AdhE spirosome structure, with a top Z-score of 48.2 and 1.8 Å R.M.S.D. (Figure S10C). In fact, the top 15 DALI results were all matches to various chains of the *Ct* AdhE complex (PDB: 8UHW), or AdhE structures from other prokaryotic organisms (Table S9). Likewise, a search using the FoldSeek server^53^ against the PDB100 database yielded an aldehyde-alcohol dehydrogenase from *E*.*coli* (PDB: 6AHC) with an expectation value of 9.9 × 10^−41^ (Figure S10D). The top 15 results from this search were all dehydrogenases - whether they be of the alcohol, aldehyde, or propionaldehyde variety (Table S10). This is consistent with the structure clustering scheme of the PDB100 database, in which models are organized based on sequence identity and coverage, and a single model is chosen to represent each cluster. Collectively, these comparisons suggested that the model built by ModelAngelo was sufficient to identify a candidate protein type (AdhE) by two independent structural search mechanisms – DALI and FoldSeek.

### Ct AdhE filaments contain a cofactor-bound ‘extended’ structure

An initial AdhE model for refinement was obtained by relaxing the structure of AdhE (PDB:8UHW) into the 4.0 □ cryoEM density map using Rosetta (Figure S11). A refined version of this model showed high similarity to published structures of the *Ct* spirosome (PDB:8UHW)^43^, with a monomer R.M.S.D. of 1.7 □, as well as to the two available *E. coli* spirosome structures in their extended forms (PDB: 7BVP, 6TQH), with monomer R.M.S.D.s of 2.2 □ and 2.93 □ respectively^42,47^. The spirosome monomer was composed of two domains, an N-terminal aldehyde dehydrogenase (ALDH) domain, and a C-terminal alcohol dehydrogenase (ADH) domain (Figure 3A-D). The ALDH domain is responsible for converting acetyl-CoA into acetaldehyde, whereas the ADH domain is responsible for further converting the reactive aldehyde to ethanol (Figure 3B). Despite the map’s relatively limited resolution, an iron atom site was identified located in the ADH domain, coordinated by three histidine residues and an aspartate residue, Asp 660, His 664, His 730 and His 744 (Figure 3D). Likewise, an NAD^+^ molecule could be modeled into semi-contiguous density within the ALDH domain (Figure 3D), with the molecule occupying a similar location to that in the *E. coli* extended spirosome structure 7BVP (Figure 3E). However, the *Ct* AdhE NAD^+^ binding cleft appeared slightly larger than that of the *E. coli* structure by about ∼200 □^3^ (Figure S12). That more open configuration is in line with the fewer contacts between the *Ct* AdhE and its bound NAD^+^, compared to its *E. coli* counterpart. It suggests that the extracellular *Ct* spirosome may adopt a state that is distinct from the previously annotated extended *E. coli* spirosome^42^, or recombinant extended apo *Ct* spirosome^43^.

**Figure 3.**
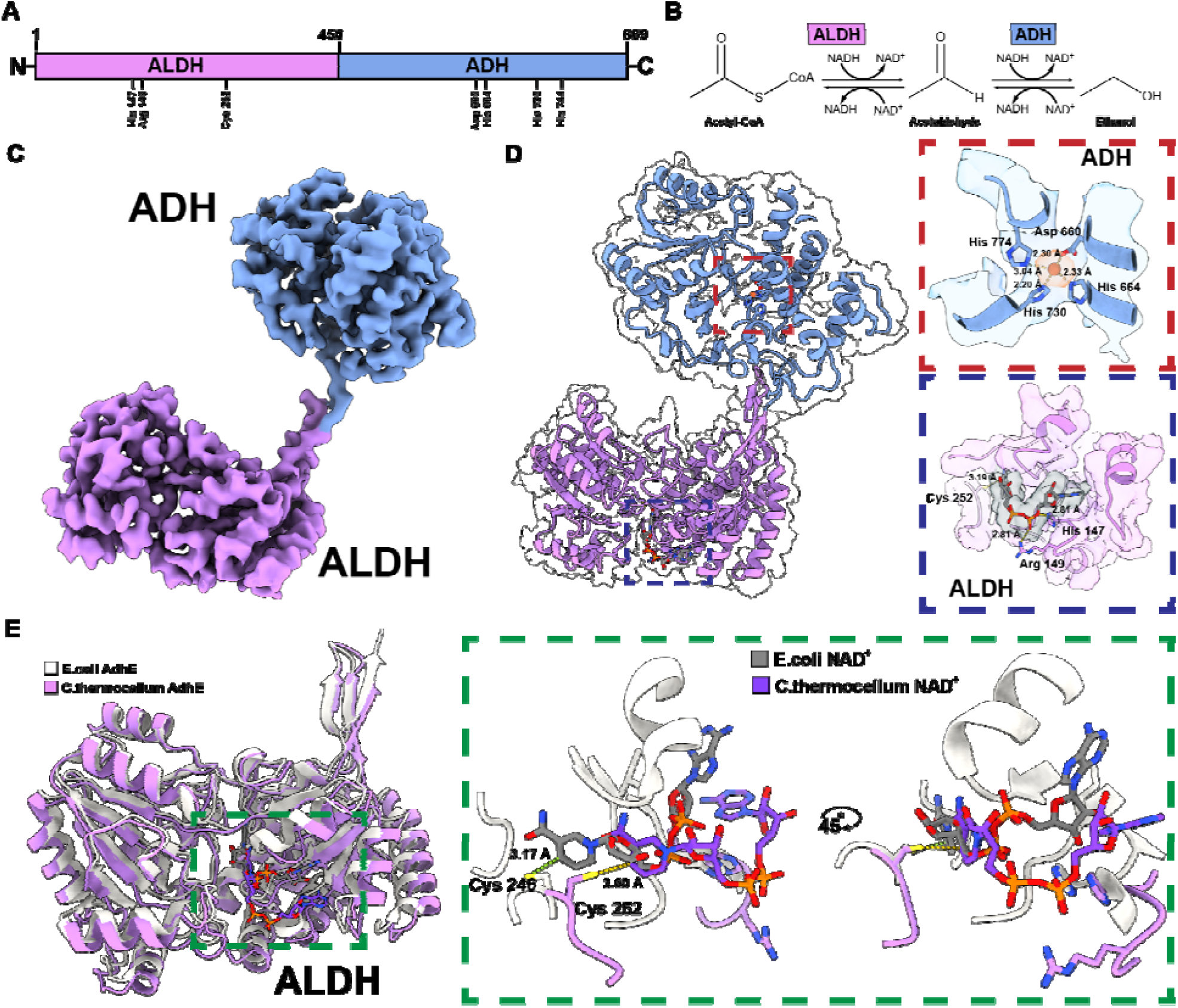
Analysis of AdhE structural features. (A) Sequence architecture of AdhE’s two-domain architecture, with important resolved interacting residues. (B) Reaction mechanism of AdhE’s conversion of acetyl-CoA to ethanol. (C) Refined map layout color coded by domain type. (D) Resolved catalytic iron atom and coordinating residues of ADH domain, top, and NAD cofactor with catalytic cysteine in ALDH domain, bottom. (E) Overlay of published *E*.*coli* AdhE structure (PDB: 7BVP) displaying similar layout but orientational differences between NAD cofactors.

### Isolated extracellular Ct spirosomes retain function

To determine whether exoproteomic *Ct* spirosomes were functional, we interrogated their ability to undergo conformational changes in response to their exposure to reactants and cofactors. We analyzed the distribution between compact and extended conformations in the assayed conditions for ethanol production as informed by nsEM and automated analysis of fibril morphologies^54^ (Figure S13). 2D classes obtained from nsEM images of each reaction condition revealed a wider conformational variety for filaments in both the forward and buffer control conditions compared to the reverse reaction condition (Figure 4A-C), potentially indicating reaction-driven changes to both the structure and biophysical properties of the isolated spirosomes. We also found that the persistence lengths of spirosomes were significantly altered in response to incubation in the various buffers, with mean lengths of 792.42 nm, 384.48 nm, and 268.84 nm for the forward reaction, reverse reaction, and purification buffers respectively (Figure 4D-F). When analyzing the total lengths of the filaments in each condition, we found no statistically significant differences (Figure 4G). We then sought to assess the ability of exoproteomic spirosomes to produce ethanol in solution when provided with acetyl-CoA and the cofactor NADH. Ethanol production measured indirectly via MTS reduction indicated enriched filament samples incubated in the forward reaction buffer dramatically increased absorption at 490 nm compared to those in the reverse reaction and purification buffers (Figure 4H). Secondly, when monitoring ethanol and acetaldehyde directly by selected ion flow tube (SIFT)-MS, we found a slight increase in ethanol concentration in spirosome-containing fractions over time for the forward reaction (Figure 4I), accompanied by a dramatic increase in acetaldehyde in both forward and reverse reactions (Figure 4J) when compared to the purification buffer control. This suggested the extracellular AdhE spirosomes were primed for function.

**Figure 4.**
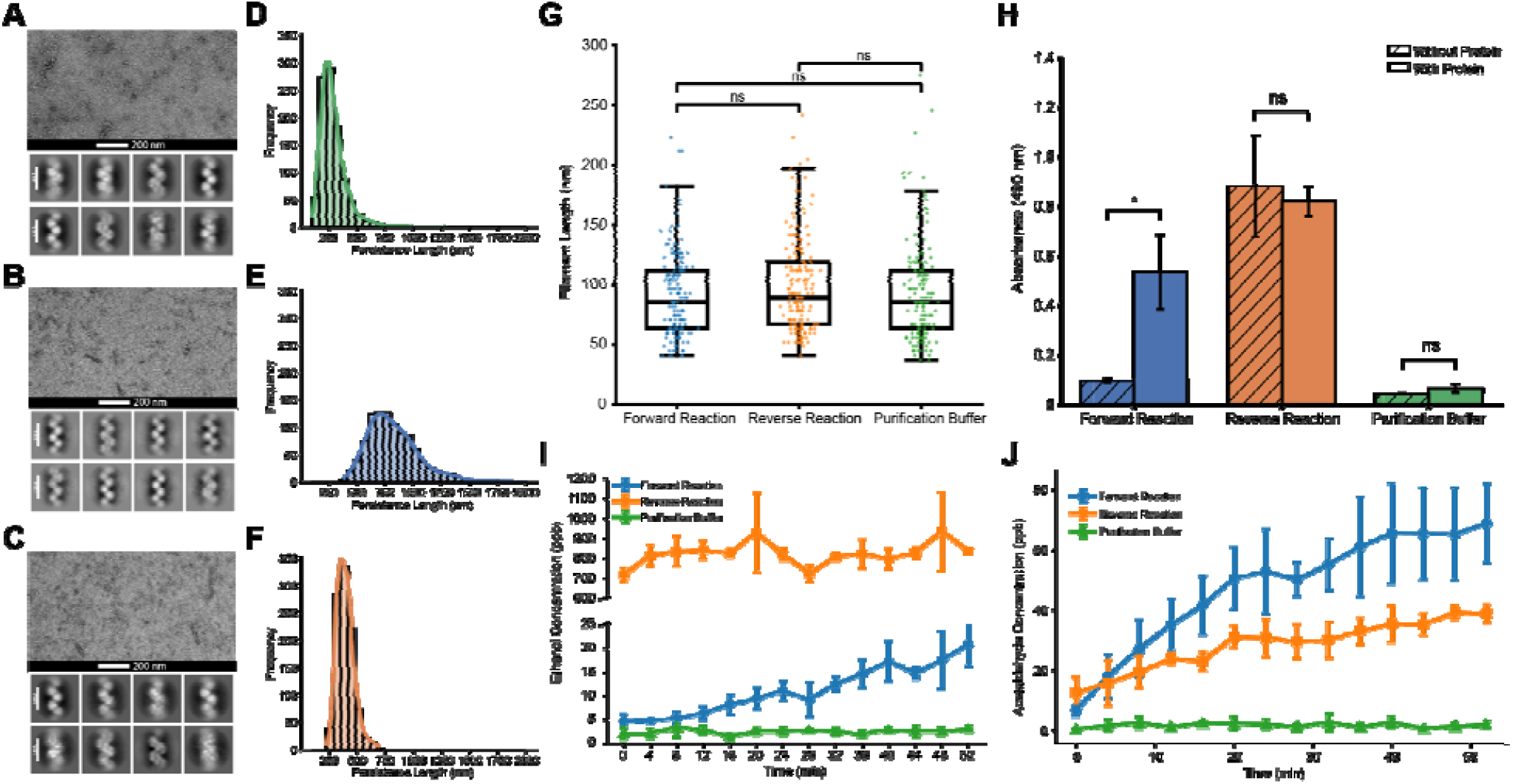
Biochemical validation of AdhE hypothesis. Conformational changes were assessed for spirosomes in the forward reaction (A), reverse reaction (B), and purification buffer (C) conditions with corresponding negative stain 2D classes illustrating diversity of conformations. Persistence lengths were analyzed using FiberApp for each condition (D-F) and show a greater persistence length compared to buffer control. (G) Overall length distributions of spirosomes were also analyzed for each condition and show no significant difference between conditions. (H) Ethanol production as assayed by proxy of MTS oxidation shows significant increase in forward reaction upon addition of enriched filaments compared to reverse reaction or purification buffer control. (I) Ethanol concentration, measured by SIFT-MS for forward, reverse and purification buffer shows slight increase for forward reaction. (J) Acetaldehyde concentration measured by SIFT-MS shows an increase for both forward and reverse reactions compared to purification buffer control.

## Discussion

Recent advances in cryoEM development have enabled high-resolution structures to be routinely obtained, with algorithmic advances allowing for the determination of multiple structures from complex mixtures of proteins^31,33,55^. Despite these advances, the accurate identification of unknown protein species from reconstructed maps alone can be challenging, particularly if the map resolution is insufficient for *de novo* model building. Mass spectrometry-based proteomics, along with software advances in model building and structure prediction^56,57^ have proven valuable for filling that void.

We leveraged the power of visual proteomics to interrogate low abundance proteins in the extracellular medium of the biomass-degrading anaerobe, *Clostridium thermocellum* (Figure S1). This complex mixture showed a wide range of proteins, including a standout filamentous complex present in high molecular weight fractions (Figure 1A,B), amidst cellulose-degrading complex components such as scaffolding proteins (Figure 1C). Although helical reconstruction of the filaments produced a map of insufficient resolution for *de novo* model building, the mass spectrometry-derived inventory of proteins within filament-containing fractions and the automated model building program ModelAngelo coupled with structural search algorithms helped guide attribution of the filament comprising protein (Figures 2, S10, Tables S3, 9,10). Results from those efforts collectively pointed to the identity of the filament protein being the aldehyde-alcohol dehydrogenase, AdhE, which is a known filamentous complex referred to as the spirosome in bacterial species (Figure 2E, S8, 9, Tables S5-8) and could be refined into the ∼4 Å filament map (Table S4). This *de novo* identification of spirosomes demonstrates the utility of visual proteomics, even when working from middling cryoEM density alone. Recent studies have further shown these tools to be accurate for poor resolution data, speaking to the robustness of these computational advances^35^.

Spirosomes from *Ct* exoproteomes are in agreement with other AdhE spirosome structures, further supporting the *de novo* identification of the complex, and raising questions about its function (Figure 2, 3, S9, S11)^42,43,58,59^. Exoproteomic spirosomes are isolated with cofactors, with evidence supporting the presence of a catalytic iron atom in the AdhE ADH domain (Figure 3D), which is coordinated by one asparagine and three histidine residues, as seen in many dehydrogenases. Evidence also supports an NAD cofactor present in the binding pocket of the ALDH domain, although limited resolution prevents the ability to distinguish between its NAD^+^ or NADH forms (Figure 3D). While the published *Ct* AdhE structure lacks density for any cofactors, NAD^+^ has been modeled into a homologous *E. coli* structure, occupying roughly the same location within the binding pocket (Figure 3E). Of note, exoproteomic spirosomes display AdhE in a slightly more elongated form than this reported extended spirosome from *E. coli*, as evidenced by the expansion in the binding pocked by almost 25% (Figure S12). This could explain the differences in residues contacting the NAD cofactor across the two structures. Importantly, the structure determined in this study represents the major population of species seen in solution, with various conformations of the filament also observed as a minor population (Figure S13).

*Ct* spirosomes in the extracellular milieu appear primed to function and can undergo conformational changes in the presence of excess acetyl-CoA or ethanol. This was demonstrated by nsEM of spirosomes incubated in buffers driving the forward or backwards biochemical reactions (Figure 3B, S14). Their incubation with acetyl-CoA and NADH induced an elongated conformation was associated with more rigid fibrils whose persistence length was approximately double that of the fibrils in the reverse reaction buffer (Figure 4D-F). Isolated spirosomes were also competent in their ability to produce ethanol (Figure 4I), but primarily generated the associated acetaldehyde intermediate with an appreciable increase in acetaldehyde concentration over time observed for both forward and reverse reactions (Figure 4J). As further conversion to ethanol may have been limited due to cofactor exhaustion or another unknown mechanism, further investigation of exoproteomic spirosome activity is warranted given the interest in *Ct*-mediated ethanol production as a potential energy substitute for fossil fuels^1,5,41,43^.

The high abundance of primed spirosomes in the exoproteome of *Ct* during anaerobic growth on cellulose raises questions about their role. While spirosomes could potentially be secreted by *Ct*, the N-terminus of AdhE lacks a secretion signal sequence, consistent with the enzyme’s role as a cytosolic protein. Further investigation of the high and medium molecular weight fractions isolated from the *Ct* media revealed an abundance of other cytosolic components including GAPDH, several porins (PorC, G, B), and various elongation factors, including EF-Tu (Figure S15, Table S11). That observation suggests that spirosomes might accumulate in the extracellular milieu from lysed cells and are enriched during harvesting and fractionation. In contrast, cellulose degrading exoproteomic components and vesicle-like objects appear more likely to be secreted. The *Ct* extracellular vesicles are particularly unusual since they appeared to be densely decorated with protein complexes and were pleomorphic. Varying in size and shape away from ideal spherical geometry, some were highly distorted into rod-like morphologies (Figure S3). Their associated proteomic components, including CbpB^46^ (Figure S4), are in line with studies suggesting that *Ct* may localize some of its cellulosome components to the surface of extracellular vesicles and that these vesicles are essential for high cellulose degradation rates^44^. As part of an ATP-binding cassette transporter system^46^, CbpB is part of a large, multi-enzyme complex that should be amenable to further investigation by cryoEM^60,61^. With several other cellulose-degrading enzymes also found to be enriched in vesicle-containing fractions, we cannot rule out the possibility that the vesicles also envelop such protein complexes (Table S2).

These findings illustrate the utility of visual proteomics at discovering extracellular complexes from native cell growth cultures. However, the current approach is limited in its ability to identify smaller or more poorly resolved assemblies. The ∼4 Å spirosome structure determined here is near the limit of what can be unambiguously identified *de novo* from visual data alone, illustrating the importance of corroborating information gleaned from mass spectra or computational predictions. Ultimately, the determination of rare complexes may require more targeted isolation procedures, and time-resolved efforts may be needed to further infer function from primed structures like the spirosome.

## Lead Contact

Further information and requests for resources and reagents should be directed the Lead Contact, Jose A. Rodriguez

## Acknowledgments

We thank Drs. Duilio Cascio and Michael Sawaya (UCLA) for their technical assistance and guidance with structure refinement. We also thank Dr. Leslie Silva (Syft Technologies, Christchurch, New Zealand) for her assistance with SIFT-MS quantification of reaction products. This work was performed in the UCLA-DOE institute for genomics and proteomics (IGP) funded by DOE award DE-FC02-02ER63421, and as a part of STROBE, an NSF Science and Technology Center through Grant DMR-1548924. Funding was also provided by NIH-NIGMS Grant R35GM128867 to J.A.R., who was also supported as a Packard Fellow, and by NIH-NIGMS Grant R35GM145286 to J.A.L. CryoEM data collection at PNNL was supported by the EMSL CryoEM Only Proposal 61815 and funded by DOE BSSD FWPl 74915 to J.E.E. Data was also collected at the Electron Imaging Center for Nanosystems (EICN) at the California NanoSystems Institute (CNSI) (NIH S10OD032459).

## Materials availability

Materials used in this study can be requested from the lead contact without restriction.

## Data availability

All data and protocols related to this study will be made available without restriction by the authors. Final model and map of AdhE are deposited under PDB code 13HE and EMDB code EMD-77065. Raw micrographs are available through EMPIAR, ID: 47488128. Raw tilt series are available through EMPIAR, ID: EMPIAR-13548. The mass spectrometry proteomics data and code used to generate figures from it have been deposited to the ProteomeXchange Consortium (http://proteomecentral.proteomexchange.com) via MassIVE partner repository with the dataset identifiers PXD078149, MSV000101762 and doi:10.25345/C5GH9BQ1G. This paper does not report original code.

## Author Contributions

J.A.R. conceptualized and directed the work. M.P.A carried out sample preparation, collected and analyzed cryoEM data, and performed model building and refinement. M.J.K. and R.R.O.L. performed mass spectrometry experiments and analysis. T.H.M. and J.E.E. performed cryoEM screening, data collection, and initial image processing at EMSL. E.H. and R.P.G. grew and processed cellular material. Y.I.G. and T.A.S. performed sample preparation. Y.I.G. and K.M.C. performed filament conformation analysis. J.A.R., R.P.G., and J.A.L. supervised the work. M.P.A and J.A.R. wrote the article with input from all authors.

## Materials and Methods

### Clostridium thermocellum culture growth

*Clostridium thermocellum* (DSM 1313; *Acetivibrio thermocellus* DSM1313) was cultured anaerobically in 250 mL serum bottles containing 100 mL of medium with vessel headspace pressurized to 10 psi in an 80:20 (v/v) mixture of N_2_CO_2_ as previously described^62^. Each 1000 mL medium of medium contained the following: 3 g Sodium Citrate Tribasic Dihydrate, 1.3 g Ammonium Sulfate, 1.56 g Potassium Phosphate Monobasic, 2.6 g MgCl_2_□6H_2_O, 0.13 g CaCl_2_□2H_2_O, 11.6 g MOPSO Sodium Salt, 4.5 g Yeast Extract (Difco), 20 μL 0.2% Resazurin solution, 10 mL of 100X Wolfe’s trace metals solution with addition of Na_2_Wo_4_□2H_2_O (0.01 g/L), 10 mL of 100X Wolfe’s vitamin solution with addition of 2-mercapto-ethane sulfonic acid (0.005 g/L)^63^ and 1 mL of a 3.5 mM solution of Ferrous Sulfate. The bottles of medium were sparged with an 80:20 (v/v) mix of N_2_CO_2_ prior to capping and sealing. Following sterilization, the medium was supplemented with 0.5 mL filter-sterilized solution of reducing agent (2.5% Na_2_S□9H_2_O, 2.5% Cysteine HCL) and 2 mL of a 1 M NaHCO_3_ solution. Following inoculation with *C. thermocellum* (2 mL OD600 ∼1.6), serum bottles were supplemented with either 2 mL of a 5% w/v solution of Avicel (Sigma Aldrich PH-101 #11365, 50 μM particle size) or 5 mL of a 20 mM cellobiose solution (Sigma Aldrich #C7252). Cells were grown to mid exponential to late log phase prior to harvest by centrifugation at low speed (500 x g) for 12 minutes at 4 °C (Fisher Scientific Centric Centrifuge). The spent cell medium was decanted and stored on ice for subsequent processing.

### Preparation of Ct exosome material

Between 400 mL to 600 mL of growth media was concentrated with 100 kDa MWCO Pierce Protein PES Concentrator tubes (Thermo Fisher Scientific) to approximately 2 mL volume. Concentrated samples were filtered with Spin-X 0.45 μm tube filters (Costar) to remove aggregates and subjected to size exclusion chromatography with a Superose 6 Increase column (Cytiva Life Sciences) using a buffer composed of 20 mM Tris pH 8.0, 150 mM NaCl, and 2 mM CaCl_2_. Fractions containing visible filaments by electron microscopy were pooled, concentrated with Vivaspin 6 300 kDa MWCO concentrators (Sartorius) to reduce the volume. Concentrated SEC fractions were placed in polycarbonate thick wall TLA-110 tubes (Beckman Coulter) and ultracentrifuged at 4 °C at 40,000 x g using an Optima Max-XP ultracentrifuge with a TLA-110 rotor (Beckman Coulter) for 1 hour to separate the filaments from the larger vesicle species. The supernatant was harvested and further concentrated to approximately 50 μL using Microcon 300 Centrifugal Biomax 300kDa concentrator tubes (Millipore Sigma) to produce an enriched filament fraction. The pellet from the ultracentrifuge run was resuspended in 10 - 12 μL of size exclusion buffer to produce an enriched vesicle fraction. Final concentrations were determined using a Nanodrop One^C^ (Thermo Fisher Scientific) using the 1 Absorbance = 1 mg/mL setting. Purified samples were stored at 4 °C until use in structural studies.

### Negative Stain EM

3 μL from the apex of each peak, diluted to 50 μg/mL was applied to freshly glow discharged Formvar/Carbon 200 mesh copper grids (Electron Microscopy Sciences) and allowed to adhere for 2 minutes. After wicking away excess liquid, grids were washed in 4 μL water, wicked, and then stained for 1.5 minutes in 4 μL of 2% uranyl acetate. Grids were screened at room temperature on Technai T12 TEM operated at 120keV (Thermo Fisher Scientific).

### Proteomics Sample Preparation

Aliquots of 10+ μg protein from each fraction were denatured in 8 M urea and reduced with 5 mM dithiothreitol (DTT) at 37 °C for 30 minutes. Reduced proteins were alkylated with 10 mM iodoacetamide (IAM) for 30 minutes in the dark at room temperature, followed by quenching with additional DTT. Protein cleanup was then performed using the SP3 protocol with SpeedBead carboxylate modified beads (Cytivia Life Sciences) and proteins were digested overnight with MS-grade trypsin at a 1:50 enzyme-to-substrate ratio (w/w). Resulting peptides were desalted using C18 stage-tips packed in-house^64^ with four layers of Empore C18 extraction disks (CDS Analytical) and dried. Dried peptides were reconstituted with 0.1% formic acid (FA) in 3% acetonitrile (ACN) in water and quantified using the Pierce Quantitative Fluorometric Peptide Assay (Thermo Fisher Scientific) prior to LC-MS/MS analysis.

### LC-MS/MS Data Acquisition

For each fraction, three technical replicate injections of 30 ng peptide were analyzed by nano-LC-MS/MS on a Q Exactive mass spectrometer (Thermo Fisher Scientific) coupled to a Dionex Ultimate 3000 RSLCnano system (Thermo Fisher Scientific). Peptides were loaded onto an Acclaim PepMap 100 C18 trap column (Thermo Fisher Scientific) at 4.5 μL/min using 0.1% FA in water as loading solvent, then separated on an Acclaim PepMap RSLC C18 analytical column (Thermo Fisher Scientific) using a 60 minute gradient of 0.1% FA in water (solvent A) and 0.1% FA in ACN (solvent B) at 300 nL/min. The gradient was as follows: 3% B for 0–5 minutes (trapping and column equilibration), 3–7% B from 5–8 minutes, 7–20% B from 8–28 minutes, 20–35% B from 28–40 min, 35–85% B from 40–42 min, held at 85% B from 42–50 minutes, then re-equilibrated to 3% B from 52–60 min. The mass spectrometer was operated in positive ion mode with a nanospray ionization (NSI) source. Source parameters were: spray voltage 1.9 kV, capillary temperature 350 °C, S-lens RF level 50. The instrument was operated in data-dependent acquisition (DDA) mode over the active gradient window (5–55 minutes). MS1 full scans were acquired in the Orbitrap over m/z 350–2,000 at a resolution of 70,000 (at m/z 200) with an automatic gain control (AGC) target of 1 × 10^6^ ions and a maximum injection time of 50 milliseconds. The top 15 most abundant precursor ions (charge states 2–6) above an intensity threshold of 4.0 × 10^4^ were selected for HCD fragmentation with a normalized collision energy (NCE) of 25. MS2 spectra were acquired in the Orbitrap at a resolution of 17,500 with an AGC target of 5 × 10^4^ ions, a maximum injection time of 50 milliseconds, and an isolation window of 2.0 m/z. Unassigned and singly charged precursors were excluded from fragmentation. Dynamic exclusion was enabled with a 40 second exclusion window. Peptide match was set to preferred and isotope exclusion was enabled.

### Proteomics Data Analysis

Raw data were processed using FragPipe v24.0^65–67^ workflow “LFQ-MBR_ with MSFragger v4.4.1 for database searching and IonQuant v1.11.20 for quantification. Spectra were searched against a target decoy *Clostridium thermocellum* proteome database (JGI, accession 2825454334) supplemented with common contaminants, using reversed sequences as decoys. Search parameters included: strict trypsin cleavage (C-terminal to K/R, no missed cleavage at KP/RP) with up to 2 missed cleavages; precursor mass tolerance of ±20 ppm; fragment mass tolerance of ±20 ppm; peptide length 7–50 residues; peptide mass range 500–5,000 Da. Carbamidomethylation of cysteine (+57.021 Da) was set as a fixed modification; oxidation of methionine (+15.995 Da) and N-terminal protein acetylation (+42.011 Da) were set as variable modifications (maximum 3 variable modifications per peptide). Isotope errors of 0, 1, and 2 were considered, and mass calibration with parameter optimization was enabled. PSM scoring and validation were performed using MSBooster (v1.4.14) and Percolator (v3.7.1). Identifications were filtered to a 1% protein-level FDR via Philosopher (v5.1.3) using the picked-protein strategy. Label-free quantification was conducted in IonQuant (v1.11.20) using match-between-runs (MBR; 0.4 minute RT tolerance) and MaxLFQ normalization. To maintain consistency with Philosopher’s primary 1% FDR filtering, IonQuant quantification thresholds were set to 100% at the peptide and protein levels. To determine relative protein composition, the intensity of an individual protein within each technical replicate was expressed as a percentage of the total summed MaxLFQ intensity for that replicate. Mean relative abundance and standard deviation were calculated across technical triplicates for each sample. Proteins were then ranked by their mean relative abundance to identify the top 10 most abundant proteins per fraction for further analysis.

### CryoEM Grid Preparation

To prepare samples for cryoEM and cryoET imaging, 3 μL of either enriched filaments or pellet vesicles at 3 mg/mL were applied to CFlat grids (CF-1.2/1.3, 300 mesh; Electron Microscopy Sciences) that had been freshly glow discharged for 20 seconds using a PELCO easiGlow. Samples were vitrified in liquid ethane using a Vitrobot Mark IV (Thermo Fisher Scientific) with a blot force of -3 N and a blot time of 4 seconds at a temperature of 12 °C and chamber humidity of 100%. Grids were screened for ice thickness and particle distribution on a Talos F200C TEM (Thermo Fisher Scientific) operated at 200KeV and equipped with a 4k x 4k Thermo Fisher Ceta 16M camera. Grids were stored in liquid nitrogen until data collection.

### Collection of CryoEM Data of Unknown Filamenteous Species

CryoEM data of enriched filaments was collected at Pacific Northwest National Labs (PNNL). Frozen grids were assembled in autogrids (Thermo Fisher Scientific) and loaded into a Titan Krios G3i (Thermo Fisher Scientific) equipped with a Gatan BioQuantum energy filter and K3 direct electron detector. Data was collected at 130 k times magnification with a beam diameter of 750 nm for a dose rate of 48 e^-^/□^2^s. Per image integration time was 1.03 seconds with a 0.0205 second frame time. The energy filter had a 20 eV slit, and the data was acquired with 4 shots per hole and a defocus range from -0.5 to -2.5 μm in -0.1 μm increments.

### CryoEM Data Processing and Model Building

CryoEM data was processed in cryoSPARC version 4.7.0^68^. Movies were aligned and summed using patch motion correction, and estimation of contrast transfer function (CTF) was performed using the patch CTF estimation job. Multiple rounds of filament tracing was carried out using 2D classes generated from the manual picking of particles from a subset of denoised micrographs with a filament diameter 0.7 □ of and a spacing of 0.2. Particles were extracted with a 512-pixel box size and no binning followed by multiple rounds of 2D classification and selected resulting in a final set of representative classes containing ∼95,000 particles. A one-volume Ab-initio model was created from the selected classes, which was used in subsequent rounds of helical refinement. A symmetry search was performed identifying the filament twist of 193.99° and rise of 59.95 □. Reference-based motion corrected and Local CTF refinement was performed on the utilized particles, followed by further helical refinement. A mask was created around a single twist of the filament and local refinement produced the final sharpened map at 4.07 □ resolution. A processing workflow can be found in Figure S7. Using the Rosetta fast relaxed^49^ model of AdhE (PDB: 8UHW) as a starting model, multiple rounds of model adjustment and refinements were carried out using the software packages Coot^69^ and Phenix^70^.

### Docking Mass Spec Hits Into Density

Using the proteins identified in the mass spec results, PDB structures or AlphaFold2-predicted models were obtained for each protein. These models were manually fitted into the density of the final unsharpened map using ChimeraX version 1.11^71^ at a map threshold of 0.4. Each model was fit multiple times in various orientations throughout the density and quality of fit was assessed by calculating the fraction of atoms that fit inside the density relative to the amount outside the density. All results were assembled into a table, ranked by best fit percentage and manually curated by protein length.

### Q-Score Analysis of Model Fits

The best fitting models were docked into the unsharpened density map in ChimeraX v1.11 and the Q-score function was performed to quantitatively determine the fit of each model for the experimental density^72^. For the top scoring model, AdhE, the Q-score analysis was repeated on a single protein chain from the experimentally determined structure (PDB 8UHW). A Rosetta fast relax procedure was subsequently performed on the entire structure followed by repeat Q-Score analysis.

### Automated Identification of Unknown Protein by Structural Features

An unsharpened version of the final density map from cryoSPARC local refinement was provided to the ModelAngelo server^51^ (https://cosmic-cryoem.org/tools/modelangelo/) without any sequence information provided as an input. After multiple rounds of automated model building, the output was loaded into Coot and individual fragments were joined into a single chain to be used as a structural search model. Fragmented chain saw searched against the entire PDB using the DALI structural search server^52^, or the PDB100 database using FoldSeek^53^ to yield a list of top matches for potential protein identity.

### Biochemical validation of AdhE Identity

For indirect measurement ethanol production, an MTS [3-(4,5-dimethylthiazol-2-yl)-5-(3-carboxymethoxyphenyl)-2-(4-sulfophenyl)-2H-tetrazolium] colorimetric assay was used to monitor NADH oxidation. Three reaction conditions were established: a forward reaction (0.1 mM NADH, 1 mM Acetyl-CoA, 1 mM FeSO_4_), a reverse reaction (1 mM NAD^+^, 1 mM Ethanol, 1 mM FeSO_4_), and a purification buffer control (20 mM Tris pH 8.0, 150 mM NaCl, 2 mM CaCl_2_). For conditions containing protein, a final concentration of 0.5 mg/mL of enriched filaments were added to each condition. Following addition of protein, MTS was added to each well at a final working concentration of 1X and allowed to incubate at room temperature. Absorbance readings were taken at 490 nm every 5 minutes for 1 hour on a VerioSkan LUX. All reactions were performed in triplicate. Ethanol and acetaldehyde production was directly measured by preparing forward, and reverse reactions as stated above along with a purification buffer control and assayed by selected ion flow tube (SIFT)-MS using a commercial SIFT Tracer i3 mass spectrometer (Syft Technologies Limited)^73,74^ operating on helium carrier gas. The SIFT-MS instrument was equipped with a multipurpose (MPS) autosampler (CTC Pal). Samples were incubated in a 6-place agitator prior to sampling of the headspace and subsequent injection into the SIFT-MS. Samples were analyzed by placing a total volume of 0.5 mL in 20 mL headspace vials incubated at 45 °C for 1 minute. A 2.5 mL aliquot of headspace was removed using a heated gas-tight syringe (120 °C) and injected into the SIFT-MS instrument’s sample inlet at 50 μL/s with a flow of make-up gas (high purity nitrogen) through a heated inlet (120 °C) to ensure that the total flow into the instrument was 25 mL/min. After sample injection, the syringe was flushed with zero air for 1 minute at 200 mL/min. A selected monitoring (SIM) method was created to measure concentrations of ethanol and acetaldehyde, using library constraints. Samples were collected using SyftAuditTracer software and analyzed in LabSyft Pro software. A dilution factor of 10.33 was used to account for the introduction of make-up gas during the headspace injection. This dilution factor was calculated with the following equation, taking into consideration the total flow, injection flow, and temperature differential between sample and syringe. The analysis time for each sample was 90 seconds and the reported concentrations are the mean of the values obtained during injection (between 35 and 50 seconds). Immediately prior to each run, enriched filaments were added to each solution at a final concentration of 0.5 mg/mL. Each reaction was performed in triplicate.

### Conformational analysis with FiberApp

Fibril tracking and analysis were performed using FiberApp^54^ to determine the contour and persistence lengths. Persistence length was determined by fitting the mean-squared end-to-end distance (MSED) to the worm-like chain model using a global pooled analysis of all fibrils. The MSED curve was weighted by the number of segment pairs at each contour length. To estimate uncertainty, fibrils were resampled with replacement (n = 1,000 bootstrap iterations), and persistence length was recalculated for each resampled dataset to generate a sampling distribution.To compare length and bootstrapped persistence length of the forward, reverse, and control samples, we implemented the Kruskal-Wallis test for non-normal distributions. For p < 0.05, pairwise comparisons were performed using Dunn’s post hoc test with Bonferroni-corrected p-values.

### Collection of CryoET Data of Enriched Vesicle Samples

CryoET data of the enriched vesicle pellet was collected at EMSL. Frozen grids were assembled into autogrids (Thermo Fisher Scientific) and loaded into a Titan Krios G3i (Thermo Fisher Scientific) equipped with a Gatan BioQuantum energy filter and K3 direct electron detector. Tilt series were collected at 33 k times magnification with a beam diameter of 3 μm and a dose rate of 2.7 e^-^/□^2^s. A dose symmetric acquisition scheme was used to collect tilt series from +60° to -60° in 3° increments. Per tilt image integration time was 1.08 seconds with a 0.09 second frame time for a total cumulative tilt series dose of ∼120 e^-^/□^2^s. The target defocus was -2.5 μm and a 20-eV slit was used.

### CryoET Data processing and segmentation

Raw frames were aligned and summed using MotionCor3^75^, aligned tilt images were restacked using in house bash scripts, followed by tilt series alignment. Aligned tilt series were input into the software package IMOD 5.1.3^76,77^ for tomogram reconstruction. Alignment was performed using the patch-tracking protocol as no fiducials were present in the sample. Tomograms were generated by weighted back-projection with 10 SIRT-like equivalents. Final tomograms were inspected by eye for quality before segmentation was performed using Dragonfly 3D World^78^ Version 2025.1 Build 14177 for Linux.

## Citations

1. Mazzoli, R. & Olson, D. G. Clostridium thermocellum: A microbial platform for high-value chemical production from lignocellulose. Adv. Appl. Microbiol. 113, 111–161 (2020).

2. Koeck, D. E., Pechtl, A., Zverlov, V. V. & Schwarz, W. H. Genomics of cellulolytic bacteria. Curr. Opin. Biotechnol. 29, 171–183 (2014).

3. Louime, C. & Uckelmann, H. Cellulosic Ethanol: Securing the Planet Future Energy Needs. Int. J. Mol. Sci. 9, 838–841 (2008).

4. Lynd, L. R., Weimer, P. J., van Zyl, W. H. & Pretorius, I. S. Microbial cellulose utilization: fundamentals and biotechnology. Microbiol. Mol. Biol. Rev. MMBR 66, 506–577, table of contents (2002).

5. Ramos, J., Valdivia, M., GarcíaLLorente, F. & Segura, A. Benefits and perspectives on the use of biofuels. Microb. Biotechnol. 9, 436–440 (2016).

6. Akinosho, H., Yee, K., Close, D. & Ragauskas, A. The emergence of Clostridium thermocellum as a high utility candidate for consolidated bioprocessing applications. Front. Chem. 2, 66 (2014).

7. Bar-On, Y. M., Phillips, R. & Milo, R. The biomass distribution on Earth. Proc. Natl. Acad. Sci. 115, 6506–6511 (2018).

8. Drula, E. et al. The carbohydrate-active enzyme database: functions and literature. Nucleic Acids Res. 50, D571–D577 (2022).

9. Bayer, E. A., Lamed, R., White, B. A. & Flint, H. J. From cellulosomes to cellulosomics. Chem. Rec. 8, 364–377 (2008).

10. Minor, C. M. et al. A genomic analysis reveals the diversity of cellulosome displaying bacteria. Front. Microbiol. 15, 1473396 (2024).

11. Lindič, N. & Vodovnik, M. Structural and functional insights into cellulosomes: masters of plant cell wall degradation. Front. Microbiol. 16, (2025).

12. Lytle, B. & Wu, J. H. D. Involvement of Both Dockerin Subdomains in Assembly of the Clostridium thermocellum Cellulosome. J. Bacteriol. 180, 6581–6585 (1998).

13. Schwarz, W. H. The cellulosome and cellulose degradation by anaerobic bacteria. Appl. Microbiol. Biotechnol. 56, 634–649 (2001).

14. Xu, Q. et al. Dramatic performance of Clostridium thermocellum explained by its wide range of cellulase modalities. Sci. Adv. 2, e1501254 (2016).

15. Yoav, S. et al. How does cellulosome composition influence deconstruction of lignocellulosic substrates in Clostridium (Ruminiclostridium) thermocellum DSM 1313? Biotechnol. Biofuels 10, 222 (2017).

16. Raman, B. et al. Impact of pretreated Switchgrass and biomass carbohydrates on Clostridium thermocellum ATCC 27405 cellulosome composition: a quantitative proteomic analysis. PloS One 4, e5271 (2009).

17. Carvalho, A. L. et al. Cellulosome assembly revealed by the crystal structure of the cohesin–dockerin complex. Proc. Natl. Acad. Sci. 100, 13809–13814 (2003).

18. Pinheiro, B. A. et al. The Clostridium cellulolyticum dockerin displays a dual binding mode for its cohesin partner. J. Biol. Chem. 283, 18422–18430 (2008).

19. Noach, I. et al. Crystal structure of a type-II cohesin module from the Bacteroides cellulosolvens cellulosome reveals novel and distinctive secondary structural elements. J. Mol. Biol. 348, 1–12 (2005).

20. Lytle, B. L., Volkman, B. F., Westler, W. M., Heckman, M. P. & Wu, J. H. Solution structure of a type I dockerin domain, a novel prokaryotic, extracellular calcium-binding domain. J. Mol. Biol. 307, 745–753 (2001).

21. García-Alvarez, B. et al. Molecular Architecture and Structural Transitions of a Clostridium thermocellum Mini-Cellulosome. J. Mol. Biol. 407, 571–580 (2011).

22. Takayesu, A. et al. Insight into the autoproteolysis mechanism of the RsgI9 anti-σ factor from Clostridium thermocellum. Proteins 92, 946–958 (2024).

23. Dumitrache, A. et al. Specialized activities and expression differences for Clostridium thermocellum biofilm and planktonic cells. Sci. Rep. 7, 43583 (2017).

24. McCafferty, C. L. et al. Integrating cellular electron microscopy with multimodal data to explore biology across space and time. Cell 187, 563–584 (2024).

25. Kelley, R. et al. Toward community-driven visual proteomics with large-scale cryoelectron tomography of Chlamydomonas reinhardtii. Mol. Cell 86, 213–230.e7 (2026).

26. Ho, C.-M. et al. Bottom-up structural proteomics: cryoEM of protein complexes enriched from the cellular milieu. Nat. Methods 17, 79–85 (2020).

27. Klykov, O. et al. Label-free visual proteomics: Coupling MS- and EM-based approaches in structural biology. Mol. Cell 82, 285–303 (2022).

28. Su, C.-C. et al. A ‘Build and Retrieve’ methodology to simultaneously solve cryo-EM structures of membrane proteins. Nat. Methods 18, 69–75 (2021).

29. Henderson, R. Avoiding the pitfalls of single particle cryo-electron microscopy: Einstein from noise. Proc. Natl. Acad. Sci. 110, 18037–18041 (2013).

30. Cheng, Y., Grigorieff, N., Penczek, P. A. & Walz, T. A Primer to Single-Particle CryoElectron Microscopy. Cell 161, 438–449 (2015).

31. Nogales, E. The development of cryo-EM into a mainstream structural biology technique. Nat. Methods 13, 24–27 (2016).

32. Cheng, Y. Single-Particle Cryo-EM at Crystallographic Resolution. Cell 161, 450–457 (2015).

33. Scheres, S. H. W. RELION: implementation of a Bayesian approach to cryo-EM structure determination. J. Struct. Biol. 180, 519–530 (2012).

34. Larance, M. & Lamond, A. I. Multidimensional proteomics for cell biology. Nat. Rev. Mol. Cell Biol. 16, 269–280 (2015).

35. Mann, M., Kulak, N. A., Nagaraj, N. & Cox, J. The coming age of complete, accurate, and ubiquitous proteomes. Mol. Cell 49, 583–590 (2013).

36. Su, C.-C. et al. High-resolution structural-omics of human liver enzymes. Cell Rep. 42, 112609 (2023).

37. Kastritis, P. L. et al. Capturing protein communities by structural proteomics in a thermophilic eukaryote. Mol. Syst. Biol. 13, 936 (2017).

38. Sae-Lee, W. et al. The protein organization of a red blood cell. Cell Rep. 40, 111103 (2022).

39. Shen, Y., Maggiolo, A. O., Zhang, T. & Warmack, R. A. CryoEM-enabled visual proteomics reveals de novo structures of oligomeric protein complexes. Structure 33, 1484–1497.e5 (2025).

40. Bayer, E. A., Chanzy, H., Lamed, R. & Shoham, Y. Cellulose, cellulases and cellulosomes. Curr. Opin. Struct. Biol. 8, 548–557 (1998).

41. Pech-Canul, A. et al. The role of AdhE on ethanol tolerance and production in Clostridium thermocellum. J. Biol. Chem. 300, 107559 (2024).

42. Kim, G. et al. Aldehyde-alcohol dehydrogenase undergoes structural transition to form extended spirosomes for substrate channeling. Commun. Biol. 3, 298 (2020).

43. Ziegler, S. J. et al. Structural characterization and dynamics of AdhE ultrastructures from Clostridium thermocellum show a containment strategy for toxic intermediates. eLife 13, RP96966 (2025).

44. Ichikawa, S. et al. Cellulosomes localise on the surface of membrane vesicles from the cellulolytic bacterium Clostridium thermocellum. FEMS Microbiol. Lett. 366, fnz145 (2019).

45. Yu, T., Xu, X., Peng, Y., Luo, Y. & Yang, K. Cell wall proteome of Clostridium thermocellum and detection of glycoproteins. Microbiol. Res. 167, 364–371 (2012).

46. Yan, F. et al. Deciphering Cellodextrin and Glucose Uptake in Clostridium thermocellum. mBio 13, e01476–22 (2022).

47. Pony, P., Rapisarda, C., Terradot, L., Marza, E. & Fronzes, R. Filamentation of the bacterial bi-functional alcohol/aldehyde dehydrogenase AdhE is essential for substrate channeling and enzymatic regulation. Nat. Commun. 11, 1426 (2020).

48. Jumper, J. et al. Highly accurate protein structure prediction with AlphaFold. Nature 596, 583–589 (2021).

49. Conway, P., Tyka, M. D., DiMaio, F., Konerding, D. E. & Baker, D. Relaxation of backbone bond geometry improves protein energy landscape modeling. Protein Sci. 23, 47–55 (2014).

50. Koehler Leman, J. et al. Macromolecular modeling and design in Rosetta: recent methods and frameworks. Nat. Methods 17, 665–680 (2020).

51. Jamali, K. et al. Automated model building and protein identification in cryo-EM maps. Nature 628, 450–457 (2024).

52. Holm, L., Laiho, A., Törönen, P. & Salgado, M. DALI shines a light on remote homologs: One hundred discoveries. Protein Sci. Publ. Protein Soc. 32, e4519 (2023).

53. van Kempen, M. et al. Fast and accurate protein structure search with Foldseek. Nat. Biotechnol. 42, 243–246 (2024).

54. Usov, I. & Mezzenga, R. FiberApp: An Open-Source Software for Tracking and Analyzing Polymers, Filaments, Biomacromolecules, and Fibrous Objects. Macromolecules 48, 1269–1280 (2015).

55. Lucas, B. A. Visualizing everything, everywhere, all at once: Cryo-EM and the new field of structureomics. Curr. Opin. Struct. Biol. 81, 102620 (2023).

56. Gao, J. et al. DomainFit: Identification of protein domains in cryo-EM maps at intermediate resolution using AlphaFold2-predicted models. Structure 32, 1248–1259.e5 (2024).

57. Bouvier, G., Bardiaux, B., Pellarin, R., Rapisarda, C. & Nilges, M. Building Protein Atomic Models from Cryo-EM Density Maps and Residue Co-Evolution. Biomolecules 12, 1290 (2022).

58. Kim, G. et al. Aldehyde-alcohol dehydrogenase forms a high-order spirosome architecture critical for its activity. Nat. Commun. 10, 4527 (2019).

59. Kessler, D., Herth, W. & Knappe, J. Ultrastructure and pyruvate formate-lyase radical quenching property of the multienzymic AdhE protein of Escherichia coli. J. jBiol. Chem. 267, 18073–18079 (1992).

60. Januliene, D. & Moeller, A. Cryo-EM of ABC transporters: an ice-cold solution to everything? FEBS Lett. 594, 3776–3789 (2020).

61. Zhang, H. et al. Cryo-EM structure of ABCG5/G8 in complex with modulating antibodies. Commun. Biol. 4, 526 (2021).

62. Wolfe, R. S. Techniques for cultivating methanogens. Methods Enzymol. 494, 1–22 (2011).

63. Sowers, K.R. & Schreir, H.J. A laboratory manual: Methanogens. Cold Spring Harbor Laboratory Press. P 474, ISBN: 0-87969-439-4 (1995).

64. Hughes, C. S. et al. Single-pot, solid-phase-enhanced sample preparation for proteomics experiments. Nat. Protoc. 14, 68–85 (2019).

65. Kong, A.T., Leprevost, F.V., Avtonomov, D.M., Mellacheruvu, D., & Nesvizhskii, A. MSFragger: ultrafast and comprehensive peptide identification in mass spectrometry-based proteomics. Nature Methods. 14, 513–520 (2017).

66. Yang, K.L., Yu, F., Teo, G.C., Li, K., Demichev, V., Ralser, M., & Nesvizhskii, A.I. MSBooster: improving peptide identification rates using deep learning-based features. Nat Commun. 14, 4539 (2023).

67. Teo, G. C., Polasky, D. A., Yu, F. & Nesvizhskii, A. I. Fast Deisotoping Algorithm and Its Implementation in the MSFragger Search Engine. J. Proteome Res. 20, 498–505 (2021).

68. Punjani, A., Zhang, H. & Fleet, D. J. Non-uniform refinement: adaptive regularization improves single-particle cryo-EM reconstruction. Nat. Methods 17, 1214–1221 (2020).

69. Emsley, P., and Cowtan, K. Coot: model-building tools for molecular graphics. Acta Crystallogr D Biol Crystallogr. 60(12), 2126–32 (2004).

70. Afonine, P. V. et al. Towards automated crystallographic structure refinement with phenix.refine. Acta Crystallogr. D Biol. Crystallogr. 68, 352–367 (2012).

71. Pettersen, E. F. et al. UCSF ChimeraX: Structure visualization for researchers, educators, and developers. Protein Sci. 30, 70–82 (2021).

72. Pintilie, G., Zhang, K., Su, Z., Li, S., Schmid, M.F., & Chiu, W. Measurement of atom resolvability in cryo-EM maps with Q-scores. Nature Methods. 17, 328–334 (2020).

73. Smith, D. & Španěl, P. Selected ion flow tube mass spectrometry (SIFT-MS) for on-line trace gas analysis. Mass Spectrom. Rev. 24, 661–700 (2005).

74. Smith, D. & Španěl, P. Recent developments and applications of selected ion flow tube mass spectrometry (SIFTLMS). Mass Spectrom. Rev. 44(2), 101–134 (2025).

75. Zheng, S. Q. et al. MotionCor2: anisotropic correction of beam-induced motion for improved cryo-electron microscopy. Nat. Methods 14, 331–332 (2017).

76. Kremer, J. R., Mastronarde, D. N. & McIntosh, J. R. Computer Visualization of Three-Dimensional Image Data Using IMOD. J. Struct. Biol. 116, 71–76 (1996).

77. Mastronarde, D. N. & Held, S. R. Automated tilt series alignment and tomographic reconstruction in IMOD. J. Struct. Biol. 197, 102–113 (2017).

78. Gendron, M. et al. Centralizing digital resources for data management, processing, and analysis for enterprise scale imaging research. Microsc. Microanal. 27, 1084–1085 (2021).

79. Nordberg, H. et al. The genome portal of the Department of Energy Joint Genome Institute. Nucleic Acids Res. 42, D26–41 (2014).

80. Schneider, C.A. et al. NIH Image to ImageJ: 25 years of image analysis. Nat Methods. 9, 671–5 (2012).

