## Supplemental tables and figures for "Visual exoproteomics of *Clostridium thermocellum* during anaerobic biomass-degradation identifies functional spirosomes"

**Supplementary Information for** "Visual exoproteomics of *Clostridium thermocellum* during anaerobic biomass-degradation identifies functional spirosomes"

**Authors:**
Matthew P. Agdanowski[1], Matthew J. Kensil[1], Trevor H. Moser[3], Ethan Humm[2], Yaneli I. Guandique[1], Kayleigh Mason-Chalmers[1], Tracy al-Set[1], Rachel R. Ogorzalek Loo[1], James E. Evans[3], Robert P. Gunsalus[2], Joseph A. Loo[1], and Jose A. Rodriguez[1*]

[1]Department of Chemistry and Biochemistry; UCLA-DOE Institute for Genomics and Proteomics; University of California, Los Angeles (UCLA), Los Angeles, CA 90095, USA
[2]Department of Microbiology Immunology and Molecular Genetics; University of California, Los Angeles (UCLA), Los Angeles, CA 90095, USA
[3]Environmental Molecular Sciences Laboratory (EMSL); Pacific Northwest National Laboratory, Richland, WA 99354, USA

### Supplementary Tables

### **Supplementary Table 1.** Label-Free Protein Quantitation for SEC Fractions 1 - 3.

| SEC Fraction | JGI Gene ID | JGI Locus Tag | Gene Product Name | Protein Description | Label-Free Quantitation (MaxLFQ) Intensity | | |
|---|---|---|---|---|---|---|---|
| | | | | | Replicate | | |
| | | | | | 1 | 2 | 3 |
| 1 | 2825455557 | Ga0373789_1226 | CbpB | Cellodextrin-binding protein B | 5.61E+09 | 5.24E+09 | 5.36E+09 |
| | 2825457206 | Ga0373789_2877 | CelS | Endoglucanase S | 1.60E+09 | 1.61E+09 | 1.67E+09 |
| | 2825456970 | Ga0373789_2641 | XynC | Xylanase C | 1.60E+09 | 1.62E+09 | 1.58E+09 |
| | 2825457081 | Ga0373789_2752 | XynZ | Xylanase Z | 1.34E+09 | 1.38E+09 | 1.37E+09 |
| | 2825456387 | Ga0373789_2058 | Rge | Rhamnogalacturonan endolyase | 9.25E+08 | 9.24E+08 | 9.76E+08 |
| | 2825454719 | Ga0373789_386 | CelG | Endoglucanase G | 7.30E+08 | 7.16E+08 | 7.25E+08 |
| | 2825456190 | Ga0373789_1861 | AdhE | Aldehyde-alcohol dehydrogenase | 7.09E+08 | 7.06E+08 | 7.26E+08 |
| | 2825454952 | Ga0373789_619 | OlpB | Outer layer protein B | 6.72E+08 | 6.87E+08 | 6.87E+08 |
| | 2825456364 | Ga0373789_2035 | CelA | Endoglucanase A | 6.03E+08 | 5.60E+08 | 6.04E+08 |
| | 2825455464 | Ga0373789_1133 | PilA | Type IV pilus assembly protein | 5.00E+08 | 5.48E+08 | 5.49E+08 |
| 2 | 2825456970 | Ga0373789_2641 | XynC | Xylanase C | 1.63E+09 | 1.72E+09 | 1.77E+09 |
| | 2825457206 | Ga0373789_2877 | CelS | Endoglucanase S | 1.53E+09 | 1.57E+09 | 1.62E+09 |
| | 2825457081 | Ga0373789_2752 | XynZ | Xylanase Z | 1.39E+09 | 1.38E+09 | 1.39E+09 |
| | 2825456364 | Ga0373789_2035 | CelA | Endoglucanase A | 9.46E+08 | 9.37E+08 | 9.57E+08 |
| | 2825456387 | Ga0373789_2058 | Rge | Rhamnogalacturonan endolyase | 8.51E+08 | 8.27E+08 | 8.92E+08 |
| | 2825454719 | Ga0373789_386 | CelG | Endoglucanase G | 6.66E+08 | 6.54E+08 | 6.71E+08 |
| | 2825456190 | Ga0373789_1861 | AdhE | Aldehyde-alcohol dehydrogenase | 5.47E+08 | 5.66E+08 | 5.66E+08 |
| | 2825455984 | Ga0373789_1653 | GH* | Uncharacterized endoglucanase | 5.20E+08 | 5.10E+08 | 5.26E+08 |
| | 2825454951 | Ga0373789_618 | CipA | Cellulosomal-scaffolding protein A | 4.85E+08 | 4.98E+08 | 5.08E+08 |
| | 2825456201 | Ga0373789_1872 | CelK | Endoglucanase K | 5.03E+08 | 4.85E+08 | 4.89E+08 |
| 3 | 2825456970 | Ga0373789_2641 | XynC | Xylanase C | 1.40E+09 | 1.49E+09 | 1.53E+09 |
| | 2825457081 | Ga0373789_2752 | XynZ | Xylanase Z | 1.28E+09 | 1.25E+09 | 1.29E+09 |
| | 2825457206 | Ga0373789_2877 | CelS | Endoglucanase S | 1.12E+09 | 1.22E+09 | 1.24E+09 |
| | 2825456190 | Ga0373789_1861 | AdhE | Aldehyde-alcohol dehydrogenase | 6.16E+08 | 6.56E+08 | 6.73E+08 |
| | 2825456387 | Ga0373789_2058 | Rge | Rhamnogalacturonan endolyase | 6.13E+08 | 5.79E+08 | 6.06E+08 |
| | 2825456364 | Ga0373789_2035 | CelA | Endoglucanase A | 5.69E+08 | 5.88E+08 | 6.08E+08 |
| | 2825454719 | Ga0373789_386 | CelG | Endoglucanase G | 4.68E+08 | 5.08E+08 | 5.07E+08 |
| | 2825455030 | Ga0373789_697 | CBM35 | Family 35 carbohydrate binding module protein | 4.20E+08 | 4.48E+08 | 4.66E+08 |
| | 2825455984 | Ga0373789_1653 | GH* | Uncharacterized endoglucanase | 4.43E+08 | 4.34E+08 | 4.40E+08 |
| | 2825454622 | Ga0373789_289 | RpsL | Small subunit ribosomal protein S12 | 4.20E+08 | 4.28E+08 | 4.31E+08 |

**Supplementary Table 2.** Label-Free Protein Quantitation for Isolated Vesicle Exosome Preparation.

| Rank | JGI Gene ID | JGI Locus Tag | Gene Product Name | Protein Description | Label-Free Quantitation (MaxLFQ) Intensity | | |
|---|---|---|---|---|---|---|---|
| | | | | | Replicate | | |
| | | | | | 1 | 2 | 3 |
| 1 | 2825455557 | Ga0373789_1226 | CbpB | Cellodextrin-binding protein B | 3.46E+09 | 3.35E+09 | 3.49E+09 |
| 2 | 2825456190 | Ga0373789_1861 | AdhE | Aldehyde-alcohol dehydrogenase | 2.25E+09 | 2.31E+09 | 2.30E+09 |
| 3 | 2825456970 | Ga0373789_2641 | XynC | Xylanase C | 1.56E+09 | 1.60E+09 | 1.59E+09 |
| 4 | 2825457081 | Ga0373789_2752 | XynZ | Xylanase Z | 1.47E+09 | 1.47E+09 | 1.48E+09 |
| 5 | 2825457206 | Ga0373789_2877 | CelS | Endoglucanase S | 1.44E+09 | 1.39E+09 | 1.42E+09 |
| 6 | 2825456387 | Ga0373789_2058 | Rge | Rhamnogalacturonan endolyase | 1.35E+09 | 1.29E+09 | 1.34E+09 |
| 7 | 2825456364 | Ga0373789_2035 | CelA | Endoglucanase A | 1.21E+09 | 1.20E+09 | 1.23E+09 |
| 8 | 2825455030 | Ga0373789_697 | CBM35 | Family 35 carbohydrate binding module protein | 5.48E+08 | 5.95E+08 | 6.03E+08 |
| 9 | 2825454719 | Ga0373789_386 | CelG | Endoglucanase G | 5.90E+08 | 5.76E+08 | 5.63E+08 |
| 10 | 2825456695 | Ga0373789_2366 | ELP* | Encapsulin/ferritin-like protein | 5.57E+08 | 5.49E+08 | 5.82E+08 |

**Supplementary Table 3.** Label-Free Protein Quantitation for Enriched Filament Exosome Preparation.

| Rank | JGI Gene ID | JGI Locus Tag | Gene Product Name | Protein Description | Label-Free Quantitation (MaxLFQ) Intensity | | |
|---|---|---|---|---|---|---|---|
| | | | | | Replicate | | |
| | | | | | 1 | 2 | 3 |
| 1 | 2825457206 | Ga0373789_2877 | CelS | Endoglucanase S | 1.79E+09 | 1.82E+09 | 1.78E+09 |
| 2 | 2825456970 | Ga0373789_2641 | XynC | Xylanase C | 1.69E+09 | 1.81E+09 | 1.84E+09 |
| 3 | 2825457081 | Ga0373789_2752 | XynZ | Xylanase Z | 1.42E+09 | 1.55E+09 | 1.53E+09 |
| 4 | 2825456190 | Ga0373789_1861 | AdhE | Aldehyde-alcohol dehydrogenase | 1.39E+09 | 1.49E+09 | 1.45E+09 |
| 5 | 2825456364 | Ga0373789_2035 | CelA | Endoglucanase A | 1.45E+09 | 1.44E+09 | 1.42E+09 |
| 6 | 2825456387 | Ga0373789_2058 | Rge | Rhamnogalacturonan endolyase | 9.51E+08 | 9.79E+08 | 9.64E+08 |
| 7 | 2825455030 | Ga0373789_697 | CBM35 | Family 35 carbohydrate binding module protein | 6.72E+08 | 7.17E+08 | 6.86E+08 |
| 8 | 2825454953 | Ga0373789_620 | Slp2 | S-layer protein 2 | 6.42E+08 | 6.35E+08 | 6.28E+08 |
| 9 | 2825454719 | Ga0373789_386 | CelG | Endoglucanase G | 5.94E+08 | 5.67E+08 | 5.95E+08 |
| 10 | 2825454951 | Ga0373789_618 | CipA | Cellulosomal-scaffolding protein A | 4.03E+08 | 4.17E+08 | 4.19E+08 |

**Table S4:** Structure determination statistics for AdhE filament.

| | |
|---|---|
| **Data collection** | |
| Magnification (kx) | 130 |
| Voltage (keV) | 300 |
| Microscope | Titan Krios |
| Camera K3 | |
| Electron exposure (e-/Å2) | 48 |
| Defocus range (μm) | -0.5 to -2.5 |
| Pixel size (Å) | 0.34 |
| **Reconstruction** | |
| Box size (pixel) | 512 |
| Final particle images (no.) | 4,863 |
| Number of particles | 94,278 |
| Helical rise (Å) | 60.06 |
| Helical twist (°) | 193.97 |
| Map resolution (Å) | 4.07 |
| FSC threshold | 0.143 |
| **Refinement** | |
| Map sharpening B factor | -74.25 |
| Model composition | |
| Non-hydrogen atoms | 40,128 |
| Protein residues | 5,166 |
| Ligands | 12 |
| R.M.S.D deviations | |
| Bond lengths (Å) | 0.002 |
| Bond angles (°) | 0.553 |
| Validation | |
| MolProbity score | 1.51 |
| Clashscore | 5.78 |
| Poor rotomers (%) | 0.00 |
| Ramachandran plot | |
| Favored (%) | 96.82 |
| Allowed (%) | 3.18 |
| Disallowed (%) | 0.00 |

**Supplementary Table 5.** Ranked list of MS protein fits docked into EM density for SEC Fraction 1.

| Locus Tag | Protein Name | Model | Total Atom Count | Atoms Outside the Contour | Atoms Inside the Contour | Fit Percentage (%) |
|---|---|---|---|---|---|---|
| Ga0373789_1861 | aldehyde-alcohol dehydrogenase | 8UHW | 30788 | 11033 | 19755 | 64.16 |
| Ga0373789_2035 | endoglucanase A | 1CEM | 3115 | 1442 | 1673 | 53.71 |
| Ga0373789_2752 | endo-1,4-beta-xylanase | 1JJF | 2250 | 1081 | 1169 | 51.96 |
| Ga0373789_1133 | type IV pilus assembly protein | AF | 1311 | 648 | 663 | 50.57 |
| Ga0373789_2752 | endo-1,4-beta-xylanase | 1JT2 | 2290 | 1179 | 1111 | 48.52 |
| Ga0373789_2035 | endoglucanase A | 1IS9 | 3582 | 1896 | 1686 | 47.07 |
| Ga0373789_1226 | sugar binding protein | 7X0I | 13193 | 7156 | 6037 | 45.76 |
| Ga0373789_1226 | sugar binding protein | AF | 3516 | 1912 | 1604 | 45.62 |
| Ga0373789_2035 | endoglucanase A | 1KWF | 3392 | 1851 | 1541 | 45.43 |
| Ga0373789_2752 | endo-1,4-beta-xylanase | 1XYZ | 5621 | 3146 | 2475 | 44.03 |
| Ga0373789_1226 | sugar binding protein | 7X0K | 13481 | 7774 | 5707 | 42.33 |
| Ga0373789_1226 | sugar binding protein | 7X0L | 13157 | 7604 | 5553 | 42.21 |
| Ga0373789_1226 | sugar binding protein | 7X0M | 7000 | 4072 | 2928 | 41.83 |
| Ga0373789_2058 | rhamnogalacturonan endolyase | AF | 6314 | 3675 | 2639 | 41.80 |
| Ga0373789_2752 | endo-1,4-beta-xylanase | AF | 6504 | 3799 | 2705 | 41.59 |
| Ga0373789_1226 | sugar binding protein | 7X0J | 13204 | 7848 | 5356 | 40.56 |
| Ga0373789_1226 | sugar binding protein | 7X0N | 13308 | 9320 | 3988 | 29.97 |
| Ga0373789_1861 | aldehyde-alcohol dehydrogenase | AF | 6736 | 5007 | 1729 | 25.67 |
| Ga0373789_2641 | endo-1,4- beta-xylanase | AF | 4908 | 3691 | 1217 | 24.80 |
| Ga0373789_2035 | endoglucanase A | AF | 3711 | 2827 | 884 | 23.82 |
| Ga0373789_386 | endoglucanase G | AF | 4463 | 3413 | 1050 | 23.53 |
| Ga0373789_2877 | cellulose 1,4-beta-cellobiosidase | AF | 5913 | 4592 | 1321 | 22.34 |
| Ga0373789_619 | anchoring scaffoldin | AF | 17467 | 13646 | 3821 | 21.88 |

**Supplementary Table 6.** Ranked list of MS protein fits docked into EM density for SEC Fraction 2.

| Locus Tag | Protein Name | Model | Total Atom Count | Atoms Outside the Contour | Atoms Inside the Contour | Fit Percentage (%) |
|---|---|---|---|---|---|---|
| Ga0373789_618 | cellulosomal-scaffolding protein A | 1ANU | 1067 | 345 | 722 | 67.67 |
| Ga0373789_1861 | aldehyde-alcohol dehydrogenase | 8UHW | 30788 | 11033 | 19755 | 64.16 |
| Ga0373789_2035 | endoglucanase A | 1CEM | 3115 | 1442 | 1673 | 53.71 |
| Ga0373789_618 | cellulosomal-scaffolding protein A | 1OHZ | 1573 | 739 | 834 | 53.02 |
| Ga0373789_2752 | endo-1,4-beta-xylanase | 1JJF | 2250 | 1081 | 1169 | 51.96 |
| Ga0373789_618 | cellulosomal-scaffolding protein A | 5G5D | 2422 | 1173 | 1249 | 51.57 |
| Ga0373789_618 | cellulosomal-scaffolding protein A | 1NBC | 2716 | 1387 | 1329 | 48.93 |
| Ga0373789_2752 | endo-1,4-beta-xylanase | 1JT2 | 2290 | 1179 | 1111 | 48.52 |
| Ga0373789_618 | cellulosomal-scaffolding protein A | 2CCL | 3674 | 1896 | 1778 | 48.39 |
| Ga0373789_618 | cellulosomal-scaffolding protein A | 2B59 | 2765 | 1430 | 1335 | 48.28 |
| Ga0373789_618 | cellulosomal-scaffolding protein A | 4B9F | 2836 | 1474 | 1362 | 48.03 |
| Ga0373789_2035 | endoglucanase A | 1IS9 | 3582 | 1896 | 1686 | 47.07 |
| Ga0373789_2035 | endoglucanase A | 1KWF | 3392 | 1851 | 1541 | 45.43 |
| Ga0373789_618 | cellulosomal-scaffolding protein A | 3KCP | 3789 | 2098 | 1691 | 44.63 |
| Ga0373789_1872 | cellulose 1,4-beta-cellobiosidase | AF | 7106 | 3977 | 3129 | 44.03 |
| Ga0373789_2752 | endo-1,4-beta-xylanase | 1XYZ | 5621 | 3146 | 2475 | 44.03 |
| Ga0373789_2058 | rhamnogalacturonan endolyase | AF | 6314 | 3675 | 2639 | 41.80 |
| Ga0373789_1653 | uncharacterized glucanase | AF | 5643 | 3294 | 2349 | 41.63 |
| Ga0373789_2752 | endo-1,4-beta-xylanase | AF | 6504 | 3799 | 2705 | 41.59 |
| Ga0373789_618 | cellulosomal-scaffolding protein A | 1AOH | 2508 | 1598 | 910 | 36.28 |
| Ga0373789_1861 | aldehyde-alcohol dehydrogenase | AF | 13204 | 7848 | 5356 | 40.56 |
| Ga0373789_2641 | endo-1,4- beta-xylanase | AF | 4908 | 3691 | 1217 | 24.80 |
| Ga0373789_2035 | endoglucanase A | AF | 3711 | 2827 | 884 | 23.82 |
| Ga0373789_386 | endoglucanase G | AF | 4463 | 3413 | 1050 | 23.53 |
| Ga0373789_2877 | cellulose 1,4-beta-cellobiosidase | AF | 5913 | 4592 | 1321 | 22.34 |
| Ga0373789_618 | cellulosomal-scaffolding protein A | AF | 13890 | 12121 | 1769 | 12.74 |

**Supplementary Table 7.** Ranked list of MS protein fits docked into EM density for SEC Fraction 3.

| Locus Tag | Protein Name | Model | Total Atom Count | Atoms Outside the Contour | Atoms Inside the Contour | Fit Percentage (%) |
|---|---|---|---|---|---|---|
| Ga0373789_1861 | aldehyde-alcohol dehydrogenase | 8UHW | 30788 | 11033 | 19755 | 64.16 |
| Ga0373789_2035 | endoglucanase A | 1CEM | 3115 | 1442 | 1673 | 53.71 |
| Ga0373789_289 | small ribosomal subunit protein uS12 | AF | 1083 | 520 | 563 | 51.99 |
| Ga0373789_2752 | endo-1,4-beta-xylanase | 1JJF | 2250 | 1081 | 1169 | 51.96 |
| Ga0373789_697 | carbohydrate binding family 6 protein | 2W47 | 1283 | 618 | 665 | 51.83 |
| Ga0373789_697 | carbohydrate binding family 6 protein | 2W1W | 2402 | 1191 | 1211 | 50.42 |
| Ga0373789_2752 | endo-1,4-beta-xylanase | 1JT2 | 2290 | 1179 | 1111 | 48.52 |
| Ga0373789_2035 | endoglucanase A | 1IS9 | 3582 | 1896 | 1686 | 47.07 |
| Ga0373789_2035 | endoglucanase A | 1KWF | 3392 | 1851 | 1541 | 45.43 |
| Ga0373789_2752 | endo-1,4-beta-xylanase | 1XYZ | 5621 | 3146 | 2475 | 44.03 |
| Ga0373789_2058 | rhamnogalacturonan endolyase | AF | 6314 | 3675 | 2639 | 41.80 |
| Ga0373789_1653 | uncharacterized glucanase | AF | 5643 | 3294 | 2349 | 41.63 |
| Ga0373789_2752 | endo-1,4-beta-xylanase | AF | 6504 | 3799 | 2705 | 41.59 |
| Ga0373789_697 | carbohydrate binding family 6 protein | AF | 6440 | 4009 | 2431 | 37.75 |
| Ga0373789_1861 | aldehyde-alcohol dehydrogenase | AF | 6736 | 5007 | 1729 | 25.67 |
| Ga0373789_2641 | endo-1,4- beta-xylanase | AF | 4908 | 3691 | 1217 | 24.80 |
| Ga0373789_2035 | endoglucanase A | AF | 3711 | 2827 | 884 | 23.82 |
| Ga0373789_386 | endoglucanase G | AF | 4463 | 3413 | 1050 | 23.53 |
| Ga0373789_2877 | cellulose 1,4-beta-cellobiosidase | AF | 5913 | 4592 | 1321 | 22.34 |

**Supplementary Table 8.** Ranked list of MS protein fits docked into EM density for enriched filament sample.

| Locus Tag | Protein Name | Model | Total Atom Count | Atoms Outside the Contour | Atoms Inside the Contour | Fit Percentage (%) |
|---|---|---|---|---|---|---|
| Ga0373789_618 | cellulosomal-scaffolding protein A | 1ANU | 1067 | 345 | 722 | 67.67 |
| Ga0373789_1861 | aldehyde-alcohol dehydrogenase | 8UHW | 30788 | 11033 | 19755 | 64.16 |
| Ga0373789_2035 | endoglucanase A | 1CEM | 3115 | 1442 | 1673 | 53.71 |
| Ga0373789_618 | cellulosomal-scaffolding protein A | 1OHZ | 1573 | 739 | 834 | 53.02 |
| Ga0373789_2752 | endo-1,4-beta-xylanase | 1JJF | 2250 | 1081 | 1169 | 51.96 |
| Ga0373789_697 | carbohydrate binding family 6 protein | 2W47 | 1283 | 618 | 665 | 51.83 |
| Ga0373789_618 | cellulosomal-scaffolding protein A | 5G5D | 2422 | 1173 | 1249 | 51.57 |
| Ga0373789_620 | cell surface glycoprotein 2 | 5G5D | 2422 | 1193 | 1229 | 50.74 |
| Ga0373789_697 | carbohydrate binding family 6 protein | 2W1W | 2402 | 1191 | 1211 | 50.42 |
| Ga0373789_618 | cellulosomal-scaffolding protein A | 1NBC | 2716 | 1387 | 1329 | 48.93 |
| Ga0373789_2752 | endo-1,4-beta-xylanase | 1JT2 | 2290 | 1179 | 1111 | 48.52 |
| Ga0373789_620 | cell surface glycoprotein 2 | 5K39 | 2777 | 1432 | 1345 | 48.43 |
| Ga0373789_618 | cellulosomal-scaffolding protein A | 2CCL | 3674 | 1896 | 1778 | 48.39 |
| Ga0373789_618 | cellulosomal-scaffolding protein A | 2B59 | 2765 | 1430 | 1335 | 48.28 |
| Ga0373789_618 | cellulosomal-scaffolding protein A | 4B9F | 2836 | 1474 | 1362 | 48.03 |
| Ga0373789_2035 | endoglucanase A | 1IS9 | 3582 | 1896 | 1686 | 47.07 |
| Ga0373789_2035 | endoglucanase A | 1KWF | 3392 | 1851 | 1541 | 45.43 |
| Ga0373789_618 | cellulosomal-scaffolding protein A | 3KCP | 3789 | 2098 | 1691 | 44.63 |
| Ga0373789_2752 | endo-1,4-beta-xylanase | 1XYZ | 5621 | 3146 | 2475 | 44.03 |
| Ga0373789_2058 | rhamnogalacturonan endolyase | AF | 6314 | 3675 | 2639 | 41.80 |
| Ga0373789_2752 | endo-1,4-beta-xylanase | AF | 6504 | 3799 | 2705 | 41.59 |
| Ga0373789_620 | cell surface glycoprotein 2 | AF | 5288 | 3228 | 2060 | 38.96 |
| Ga0373789_697 | carbohydrate binding family 6 protein | AF | 6440 | 4009 | 2431 | 37.75 |
| Ga0373789_618 | cellulosomal-scaffolding protein A | 1AOH | 2508 | 1598 | 910 | 36.28 |
| Ga0373789_1861 | aldehyde-alcohol dehydrogenase | AF | 6736 | 5007 | 1729 | 25.67 |

| Locus Tag | Protein Name | Model | Total Atom Count | Atoms Outside the Contour | Atoms Inside the Contour | Fit Percentage (%) |
|---|---|---|---|---|---|---|
| Ga0373789_618 | cellulosomal-scaffolding protein A | 1ANU | 1067 | 345 | 722 | 67.67 |
| Ga0373789_1861 | aldehyde-alcohol dehydrogenase | 8UHW | 30788 | 11033 | 19755 | 64.16 |
| Ga0373789_2641 | endo-1,4- beta-xylanase | AF | 4908 | 3691 | 1217 | 24.80 |
| Ga0373789_2035 | endoglucanase A | AF | 3711 | 2827 | 884 | 23.82 |
| Ga0373789_386 | endoglucanase G | AF | 4463 | 3413 | 1050 | 23.53 |
| Ga0373789_2877 | cellulose 1,4-beta-cellobiosidase | AF | 5913 | 4592 | 1321 | 22.34 |
| Ga0373789_618 | cellulosomal-scaffolding protein A | AF | 13890 | 12121 | 1769 | 12.74 |

**Supplementary Table 9.** Top 15 potential identities from DALI structural search against the entire PDB using ModelAngelo fragment prediction model as search query.

| **DALI Full PDB Top 15 Protein Results** | | | | | | |
|---|---|---|---|---|---|---|
| **Chain Identifier** | **Z-Score** | **R.M.S.D. (Å)** | **Alignment Length** | **Number of Residues** | **Percent Identity (%)** | **PDB Description** |
| 8UHW_E | 48.2 | 1.8 | 797 | 859 | 46 | Aldehyde-Alcohol Dehydrogenase |
| 8UHW_D | 48.2 | 1.7 | 797 | 859 | 46 | Aldehyde-Alcohol Dehydrogenase |
| 8UHW_C | 47.9 | 1.7 | 797 | 859 | 46 | Aldehyde-Alcohol Dehydrogenase |
| 6TQM_C | 45.5 | 1.6 | 396 | 420 | 32 | Aldehyde-Alcohol Dehydrogenase |
| 6TQM_F | 45.3 | 1.6 | 396 | 420 | 32 | Aldehyde-Alcohol Dehydrogenase |
| 7BVP_F | 45.1 | 2.2 | 797 | 869 | 35 | Aldehyde-Alcohol Dehydrogenase |
| 7BVP_D | 45.1 | 2.2 | 797 | 869 | 35 | Aldehyde-Alcohol Dehydrogenase |
| 7BVP_B | 45.1 | 2.2 | 797 | 869 | 35 | Aldehyde-Alcohol Dehydrogenase |
| 7BVP_A | 45.1 | 2.2 | 797 | 869 | 35 | Aldehyde-Alcohol Dehydrogenase |
| 7BVP_E | 45.1 | 2.2 | 797 | 869 | 35 | Aldehyde-Alcohol Dehydrogenase |
| 7BVP_C | 45.1 | 2.2 | 797 | 869 | 35 | Aldehyde-Alcohol Dehydrogenase |
| 6AHC_H | 45.1 | 1.7 | 395 | 419 | 33 | Aldehyde-Alcohol Dehydrogenase |
| 6AHC_G | 45.1 | 4.2 | 457 | 869 | 30 | Aldehyde-Alcohol Dehydrogenase |
| 6AHC_A | 45.0 | 1.7 | 394 | 419 | 33 | Aldehyde-Alcohol Dehydrogenase |
| 6AHC_C | 44.6 | 4.4 | 458 | 869 | 30 | Aldehyde-Alcohol Dehydrogenase |

**Supplementary Table 10.** Top 15 potential identities from FoldSeek structural search of the PDB100 database using ModelAngelo fragment prediction model as search query.

| FoldSeek PDB100 Top 15 Protein Results | | | | | |
|---|---|---|---|---|---|
| **Target** | **Description** | **Scientific Name** | **Probability** | **Sequence Identity (%)** | **E-Value** |
| 6AHC_Assembly1_B | Cryo-EM structure of aldehyde-alcohol dehydrogenase | *Escherichia coli* K-12 | 1.00 | 33.6 | 9.90E-41 |
| 7DAG_Assembly1_A | Vibrio cholera aldehyde-alcohol dehydrogenase | *Vibrio cholerae* | 1.00 | 30.2 | 2.69E-36 |
| 7DAG_Assembly1_H | Vibrio cholera aldehyde-alcohol dehydrogenase | *Vibrio cholerae* | 1.00 | 29.4 | 2.39E-28 |
| 3MY7_Assembly2_B | Crystal structure of the ACDH domain of an alcohol dehydrogenase | *Vibrio parahaemolyticus* RMID 2210633 | 1.00 | 29.8 | 1.98E+27 |
| 3MY7_Assembly2_C | Crystal structure of the ACDH domain of an alcohol dehydrogenase | *Vibrio parahaemolyticus RMID 2210633* | 1.00 | 30.2 | 7.03E-27 |
| 3K9D_Assembly2_D | Crystal structure of probable aldehyde dehydrogenase | *Listeria monocytogenes* | 1.00 | 26.3 | 1.14E-26 |
| 3MY7_Assembly1_D | Crystal structure of ACDH domain of an alcohol dehydrogenase | *Vibrio parahaemolyticus* RMID 2210633 | 1.00 | 30 | 1.74E-26 |
| 5J7I_Assembly1_A | Crystal structure of acetylating aldehyde dehydrogenase | *Parageobacillus thermoglucosidasius* C56-YS93 | 1.00 | 29.4 | 5.16E-265 |
| 3MY7_Assembly1_A | Crystal structure of the ACDH domain of an alcohol dehydrogenase | *Vibrio parahaemolyticus* RMID 2210633 | 1.00 | 29.5 | 6.19E-26 |
| 5J78_Assembly1_D | Crystal structure of acetylating aldehyde dehydrogenase | *Parageobacillus thermoglucosidasius* | 1.00 | 29 | 9.45E-26 |
| 8CEJ_Assembly1_A | Crystal structure of succinyl-CoA reductase | *Clostridium kluyveri* | 1.00 | 23.2 | 9.25E-23 |
| 4C3S_Assembly1_A-2 | Crystal structure of propionaldehyde dehydrogenase | *Lachnoclostridium phytofermantans* | 1.00 | 20.3 | 4.19E-19 |
| 5DBV_Assembly1_A | Crystal structure of a C269A mutant of propionaldehyde dehydrogenase | *Lachnoclostridium phytofermantans* ISDg | 1.00 | 20.3 | 1.08E-18 |
| 5DRU_Assembly1_A | Crystal structure of a H387A mutant of propionaldehyde dehydrogenase | *Lachnoclostridium phytofermantans* | 1.00 | 21.1 | 2.36E-18 |
| 5JFM_Assembly2_F | Crystal structure of propionaldehyde dehydrogenase | *Rhodopseudomonas palustris* BisB18 | 1.00 | 16.1 | 7.44E-18 |

**Supplemental Table 11.** Likely cytoplasmic protein markers from lower molecular weight SEC fractions.

| Locus Tag | Protein Description | Label-Free Quantitation (MaxLFQ) Intensity | | | COG Category | COG Function |
|---|---|---|---|---|---|---|
| | | Replicate 1 | Replicate 2 | Replicate 3 | | |
| Ga0373789_2754 | NADH-dependent peroxiredoxin subunit C | 1557103740 | 1514801150 | 1451347460 | V | Defense Mechanisms |
| Ga0373789_14 | cellulose binding protein with CBM3 domain | 902213950 | 923222910 | 913517250 | N/A | No assigned COG |
| Ga0373789_3200 | pyruvate ferredoxin oxidoreductase gamma subunit | 768655740 | 745024640 | 754794370 | C | Energy production and conversion |
| Ga0373789_3202 | pyruvate ferredoxin oxidoreductase alpha subunit | 732353470 | 743524670 | 744456640 | C | Energy production and conversion |
| Ga0373789_697 | lysophospholipase L1-like esterase | 624101760 | 665878590 | 655036030 | DI | Cell cycle control, cell division, chromosomes partitioning |
| Ga0373789_1242 | elongation factor Ts | 655209220 | 651351870 | 621565500 | J | Translation, ribosomal structure and biogenesis |
| Ga0373789_292 | elongation factor Tu | 551026300 | 555967740 | 546341120 | J | Translation, ribosomal structure and biogenesis |
| Ga0373789_2181 | triosephosphate isomerase (TIM) | 552510400 | 550327100 | 522733344 | G | Carbohydrate transport and metabolism |
| Ga0373789_2994 | xylan 1,4-beta-xylosidase | 521996128 | 467181440 | 492725536 | G | Carbohydrate transport and metabolism |
| Ga0373789_3203 | pyruvate ferredoxin oxidoreductase beta subunit | 458415808 | 443493952 | 441260032 | C | Energy production and conversion |
| Ga0373789_1557 | small subunit ribosomal protein S1/4-hydroxy-3-methylbut-2-en-1-yl diphosphate reductase | 437327776 | 450355488 | 414913184 | J | Translation, ribosomal structure and biogenesis |
| Ga0373789_2183 | glyceraldehyde 3-phosphate dehydrogenase (phosphorylating) | 344452992 | 383203328 | 352363008 | G | Carbohydrate transport and metabolism |
| Ga0373789_2096 | glucose-6-phosphate isomerase | 330184384 | 297783456 | 296983744 | G | Carbohydrate transport and metabolism |
| Ga0373789_2029 | cellobiose phosphorylase | 280320384 | 295723840 | 292338720 | G | Carbohydrate transport and metabolism |
| Ga0373789_1910 | glutamate dehydrogenase (NADP+) | 247474064 | 253115392 | 250904992 | E | Amino acid transport and metabolism |
| Ga0373789_2178 | enolase 1/2/3 | 244499248 | 248882208 | 252707568 | G | Carbohydrate transport and metabolism |
| Ga0373789_525 | cellobiose phosphorylase | 225246752 | 224471008 | 218516016 | G | Carbohydrate transport and metabolism |
| Ga0373789_1945 | malate dehydrogenase (oxaloacetate-decarboxylating) | 172109120 | 181266160 | 177626176 | C | Energy production and conversion |
| Ga0373789_276 | dihydroxy-acid dehydratase | 174769744 | 174157472 | 176046064 | EG | Amino acid transport and metabolism |
| Ga0373789_301 | trigger factor | 171128704 | 168148656 | 170451136 | O | Posttranslational modification, protein turnover, chaperones |
| Ga0373789_2549 | NifU-like protein involved in Fe-S cluster formation | 165456080 | 170494112 | 153511360 | O | Posttranslational modification, protein turnover, chaperones |
| Ga0373789_1944 | L-lactate dehydrogenase | 167878720 | 156640592 | 157018992 | C | Energy production and conversion |

| | | | | | | |
|---|---|---|---|---|---|---|
| Ga0373789_422 | large subunit ribosomal protein L3 | 25835694 | 115491688 | 318794112 | J | Translation, ribosomal structure and biogenesis |
| Ga0373789_1292 | SpoIID/LytB domain protein | 147420928 | 153487664 | 156983760 | DM | Cell cycle control, cell division, chromosomes partitioning |
| Ga0373789_170 | rod shape-determining protein MreB and related proteins | 0 | 205287440 | 213402640 | DZ | Cell cycle control, cell division, chromosomes partitioning |
| Ga0373789_676 | pyruvate-ferredoxin/flavodoxin oxidoreductase | 132898152 | 154188992 | 122943120 | F | Nucleotide transport and metabolism |
| Ga0373789_1478 | isoleucyl-tRNA synthetase | 125020480 | 140384512 | 137428320 | J | Translation, ribosomal structure and biogenesis |
| Ga0373789_1942 | ATP-dependent phosphofructokinase / diphosphate-dependent phosphofructokinase | 128191048 | 132302912 | 129932624 | G | Carbohydrate transport and metabolism |
| Ga0373789_888 | aspartate kinase | 135751552 | 124047536 | 124121160 | E | Amino acid transport and metabolism |
| Ga0373789_2548 | GGGtGRT protein | 124952880 | 128339056 | 113948784 | | No assigned COG |
| Ga0373789_2750 | mannose-1-phosphate guanylyltransferase / phosphomannomutase | 113055696 | 122605232 | 105060768 | G | Carbohydrate transport and metabolism |
| Ga0373789_563 | 2-methylcitrate synthase | 103325296 | 103218648 | 105423768 | C | Energy production and conversion |
| Ga0373789_680 | long-chain acyl-CoA synthetase | 102539160 | 103157616 | 101981328 | I | Lipid transport and metabolism |
| Ga0373789_302 | ATP-dependent Clp protease, protease subunit | 90133424 | 106685808 | 101411224 | O | Posttranslational modification, protein turnover, chaperones |
| Ga0373789_2019 | isocitrate dehydrogenase | 101893144 | 97892464 | 92212688 | C | Energy production and conversion |
| Ga0373789_1525 | adenylosuccinate lyase | 97320400 | 98712976 | 93475968 | F | Nucleotide transport and metabolism |
| Ga0373789_65 | 2-isopropylmalate synthase | 95444656 | 94307296 | 94061304 | E | Amino acid transport and metabolism |
| Ga0373789_1030 | phosphoribosylaminoimidazolecarboxamide formyltransferase / IMP cyclohydrolase | 95744600 | 93284056 | 91069296 | F | Nucleotide transport and metabolism |
| Ga0373789_64 | ketol-acid reductoisomerase | 93718592 | 89587432 | 91110072 | EH | Amino acid transport and metabolism |
| Ga0373789_2366 | uncharacterized protein | 89515304 | 97649712 | 84060768 | P | Inorganic ion transport and metabolism |
| Ga0373789_1398 | 2-oxoglutarate/2-oxoacid ferredoxin oxidoreductase subunit beta | 84670800 | 97228480 | 86162456 | C | Energy production and conversion |
| Ga0373789_1666 | indolepyruvate ferredoxin oxidoreductase, alpha subunit | 92319624 | 91623360 | 83689176 | C | Energy production and conversion |
| Ga0373789_476 | prolyl-tRNA synthetase | 86903336 | 89749552 | 88133032 | J | Translation, ribosomal structure and biogenesis |
| Ga0373789_2753 | NADH-dependent peroxiredoxin subunit F | 87295832 | 79933392 | 81865576 | V | Defense Mechanisms |
| Ga0373789_571 | D-3-phosphoglycerate dehydrogenase / 2-oxoglutarate reductase | 83799368 | 80531248 | 82644288 | H | Coenzyme transport and metabolism |
| Ga0373789_462 | glycosyl hydrolase family 18 (putative chitinase) | 77447192 | 87819968 | 81442824 | | No assigned COG |

| | | | | | | |
|---|---|---|---|---|---|---|
| Ga0373789_1397 | 2-oxoglutarate/2-oxoacid ferredoxin oxidoreductase subunit alpha | 82866960 | 75101064 | 81518544 | C | Energy production and conversion |
| Ga0373789_2222 | alcohol dehydrogenase | 78633096 | 80345376 | 80320024 | C | Energy production and conversion |
| Ga0373789_406 | chaperonin GroEL | 77889264 | 74495760 | 77848272 | O | Posttranslational modification, protein turnover, chaperones |
| Ga0373789_1271 | stage V sporulation protein D (sporulation-specific penicillin-binding protein) | 79888968 | 78448296 | 70673032 | DM | Cell cycle control, cell division, chromosome partitioning |
| Ga0373789_1971 | valyl-tRNA synthetase | 73903440 | 77394744 | 75874824 | J | Translation, ribosomal structure and biogenesis |
| Ga0373789_2884 | methionyl-tRNA synthetase | 76602992 | 76147200 | 73778008 | J | Translation, ribosomal structure and biogenesis |
| Ga0373789_2543 | HlyD family secretion protein | 74678456 | 73730432 | 72420520 | MV | Cell wall/membrane/envelope biogenesis |
| Ga0373789_2141 | argininosuccinate synthase | 70681648 | 69542776 | 64012244 | E | Amino acid transport and metabolism |
| Ga0373789_1027 | amidophosphoribosyltransferase | 63106328 | 64493704 | 73078368 | F | Nucleotide transport and metabolism |
| Ga0373789_2645 | O-acetylhomoserine (thiol)-lyase | 64057536 | 66465844 | 69055312 | E | Amino acid transport and metabolism |
| Ga0373789_1138 | hypothetical protein | 89448064 | 57642816 | 43873384 | | No assigned COG |
| Ga0373789_1218 | acetate kinase | 67522304 | 62732752 | 59296344 | C | Energy production and conversion |
| Ga0373789_1854 | NADH-quinone oxidoreductase subunit G | 40337816 | 101392368 | 46060052 | C | Energy production and conversion |
| Ga0373789_1048 | threonyl-tRNA synthetase | 63483744 | 63701456 | 60052984 | J | Translation, ribosomal structure and biogenesis |
| Ga0373789_3201 | pyruvate ferredoxin oxidoreductase delta subunit | 61676076 | 59341032 | 63172236 | C | Energy production and conversion |
| Ga0373789_1241 | small subunit ribosomal protein S2 | 62845884 | 60022264 | 58894660 | J | Translation, ribosomal structure and biogenesis |
| Ga0373789_1723 | eukaryotic-like serine/threonine-protein kinase | 60859528 | 57992124 | 62165556 | T | Signal transduction mechanisms |
| Ga0373789_1399 | 2-oxoglutarate ferredoxin oxidoreductase subunit gamma | 57843440 | 61843008 | 59410260 | C | Energy production and conversion |
| Ga0373789_291 | elongation factor G | 59573284 | 57386292 | 58289244 | J | Translation, ribosomal structure and biogenesis |
| Ga0373789_989 | serine protease Do | 56978124 | 53814328 | 55655732 | O | Posttranslational modification, protein turnover, chaperones |
| Ga0373789_1909 | GMP synthase (glutamine-hydrolysing) | 53273696 | 54557808 | 53801176 | F | Nucleotide transport and metabolism |
| Ga0373789_1780 | formate C-acetyltransferase | 52561224 | 52681580 | 50482852 | C | Energy production and conversion |
| Ga0373789_181 | ribose-phosphate pyrophosphokinase | 53985984 | 51621948 | 49524556 | EF | Amino acid transport and metabolism |
| Ga0373789_1855 | NADH-quinone oxidoreductase subunit F | 52266160 | 51368312 | 50879724 | C | Energy production and conversion |
| Ga0373789_2619 | hypothetical protein | 47502016 | 56894776 | 50033752 | | No assigned COG |
| Ga0373789_1861 | acetaldehyde dehydrogenase / alcohol dehydrogenase | 50582028 | 51704144 | 47119904 | I | Lipid transport and metabolism |

| | | | | | | |
|---|---|---|---|---|---|---|
| Ga0373789_1860 | RIO-like serine/threonine protein kinase | 50143200 | 45247396 | 53098816 | T | Signal transduction mechanisms |
| Ga0373789_3189 | seryl-tRNA synthetase | 42544280 | 50236068 | 52459660 | J | Translation, ribosomal structure and biogenesis |
| Ga0373789_1546 | tyrosyl-tRNA synthetase | 48094380 | 43071724 | 53406696 | J | Translation, ribosomal structure and biogenesis |

### Supplementary Figures

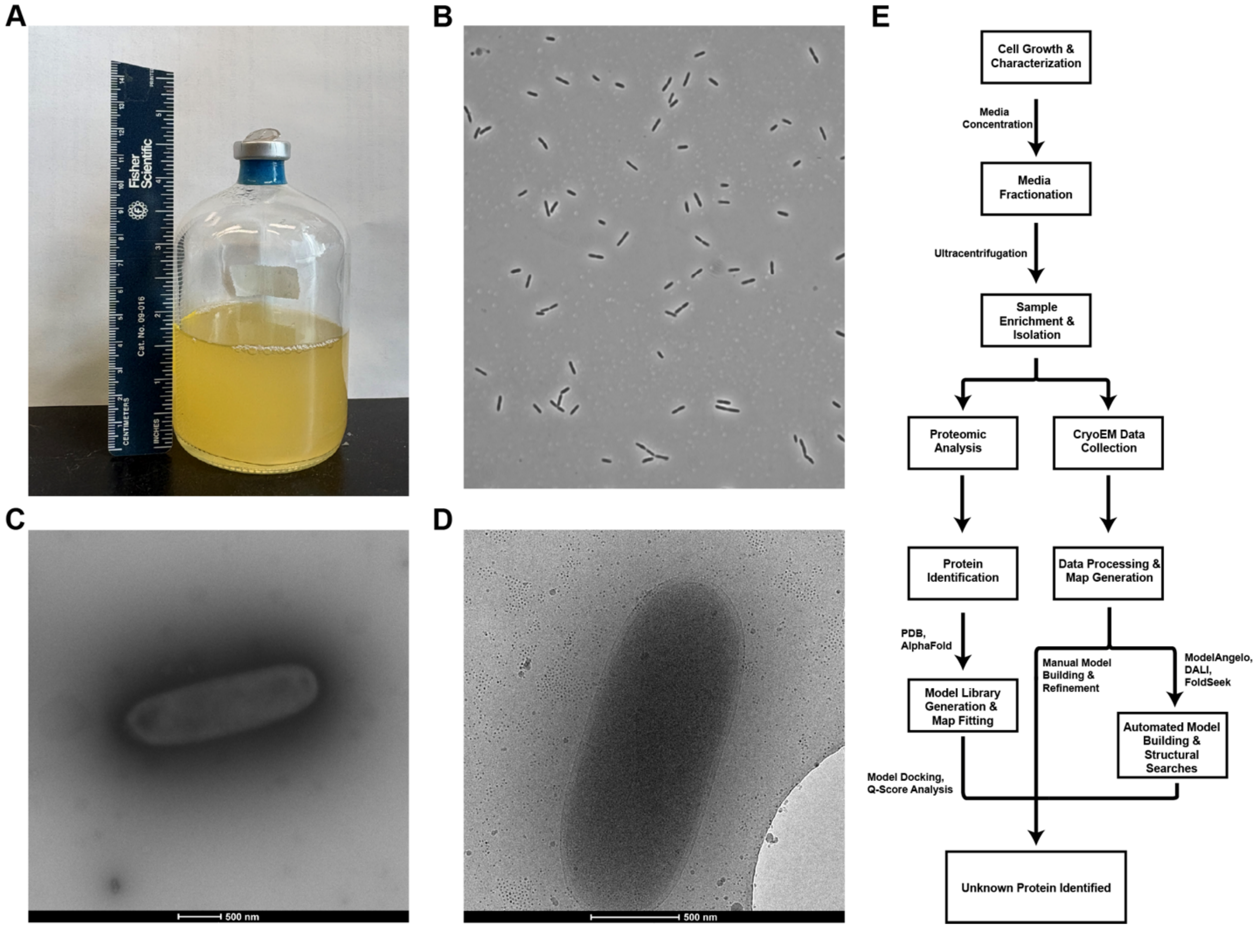


**Supplemental Figure 1**. *C. thermocellum* growth and morphological characterization. (A) Image of *Ct* cellobiose anaerobic culture. (B) Phase contrast image of *Ct* cells. (C) Negative stain electron micrograph of *Ct* cells. (D) cryoEM micrograph of a whole *Ct* cell. In all observations, *Ct* cells display characteristic rod-like morphology, indicating healthy cell growth. (E) Visual exoproteomics workflow established to characterize *C. thermocellum* complexes during anaerobic growth on cellobiose.

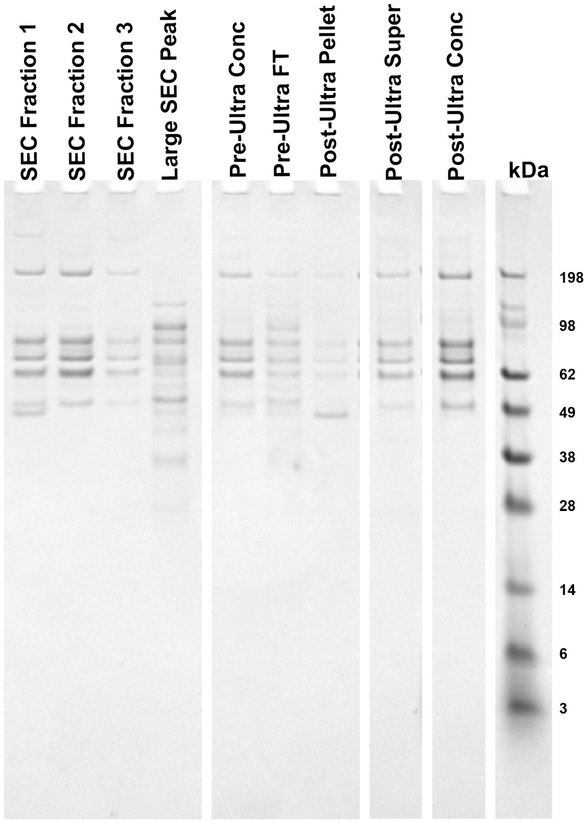


**Supplemental Figure 2.** SDS-PAGE of exosome preparation steps. SEC Fractions 1-3 correspond to high molecular weight peaks that pooled for further prep (Lanes 1-3). Fractions were further concentrated (Lane 5) before being subjected to ultracentrifugation. The pellet following ultracentrifugation was resuspended to create an isolated vesicle fraction (Lane 7), and the supernatant was further concentrated to produce an enriched filament fraction (Lane 9) used in further structural and biochemical studies.

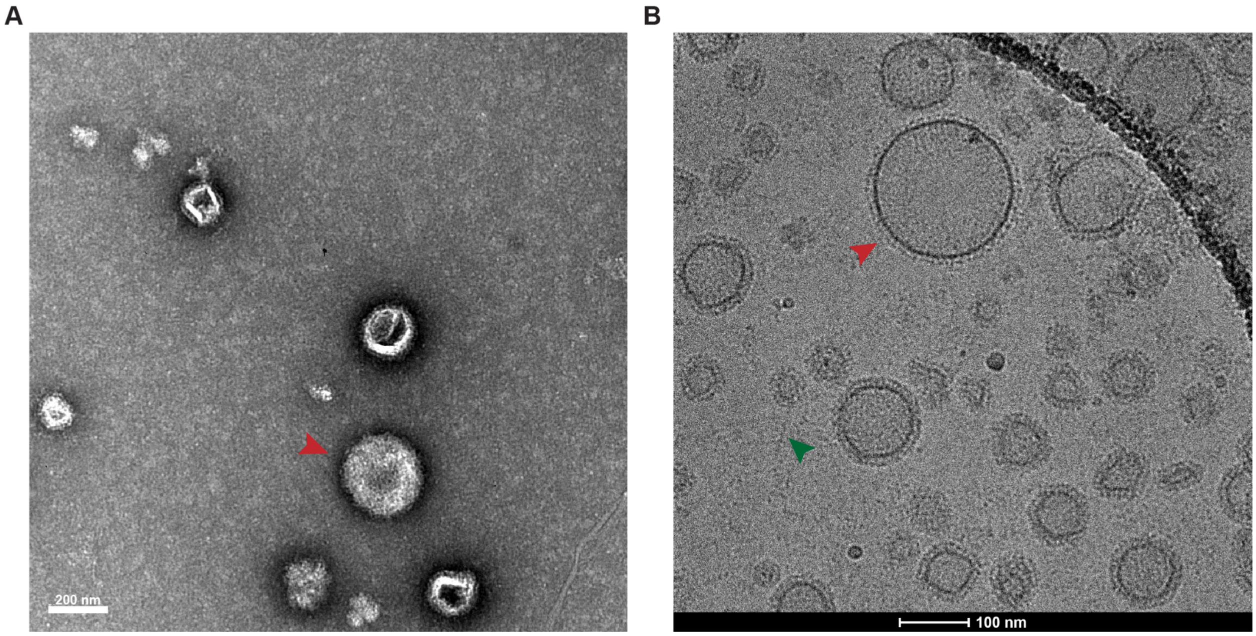


**Supplemental Figure 3.** Characterization of vesicle species by EM. Ultracentrifugation of fractionated media contains vesicle species of varying sizes in the pellet fraction. (A)Negative stain electron microscopy, and (B) cryoEM reveal morphological differences as well as clear proteinaceous decorations. Example vesicles are denoted with red arrows, filaments are denoted by green arrows.

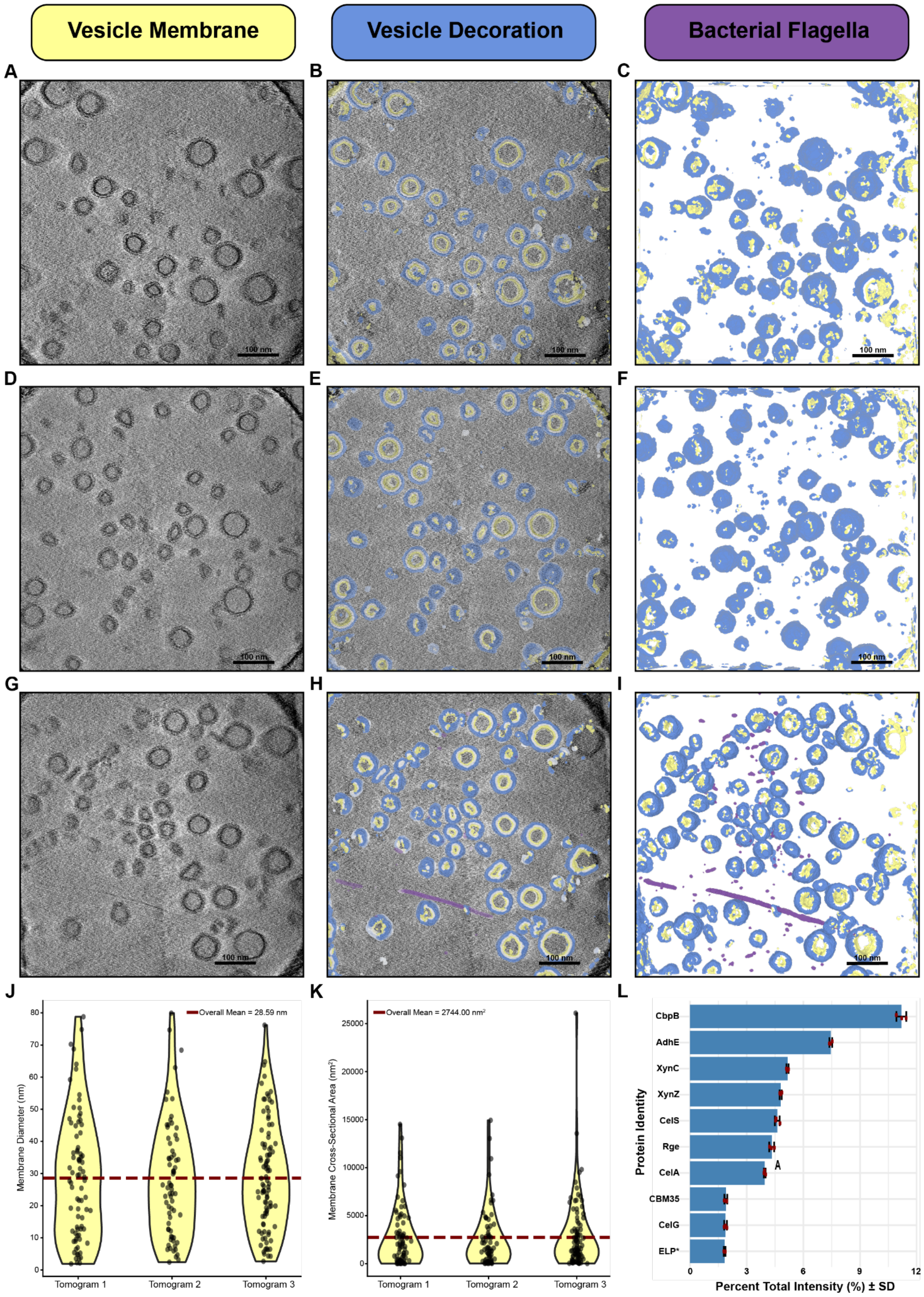


**Supplemental Figure 4.** Investigation of vesicle-like species. (A, D, G) Tomogram slices (20 frames summed) depicting the size and morphology distribution of isolated vesicles and their decoration. (overlay: B, E, H isolated: C, F, I) Segmentation of individual species from reconstructed tomograms of isolated vesicles showing presence of decorated vesicles, bacteria flagella. (J) Distribution of segmented membrane diameters across all three tomograms. (K) Distribution of vesicle membrane cross-sectional area for each tomogram. (L) Bottom-up proteomics of isolated vesicle fractions reveal sugar-binding protein, CbpB as the most abundant species.

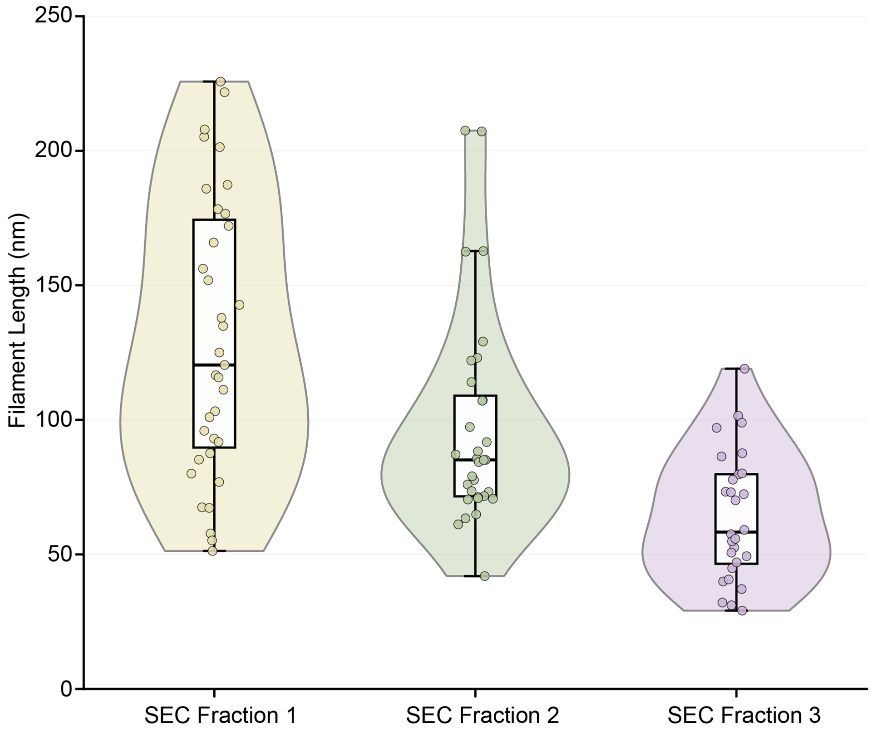


**Supplemental Figure 5.** Filament length distribution in each of the high molecular weight SEC fractions. Lengths of filaments decrease as a function of elution volume in the size exclusion column.

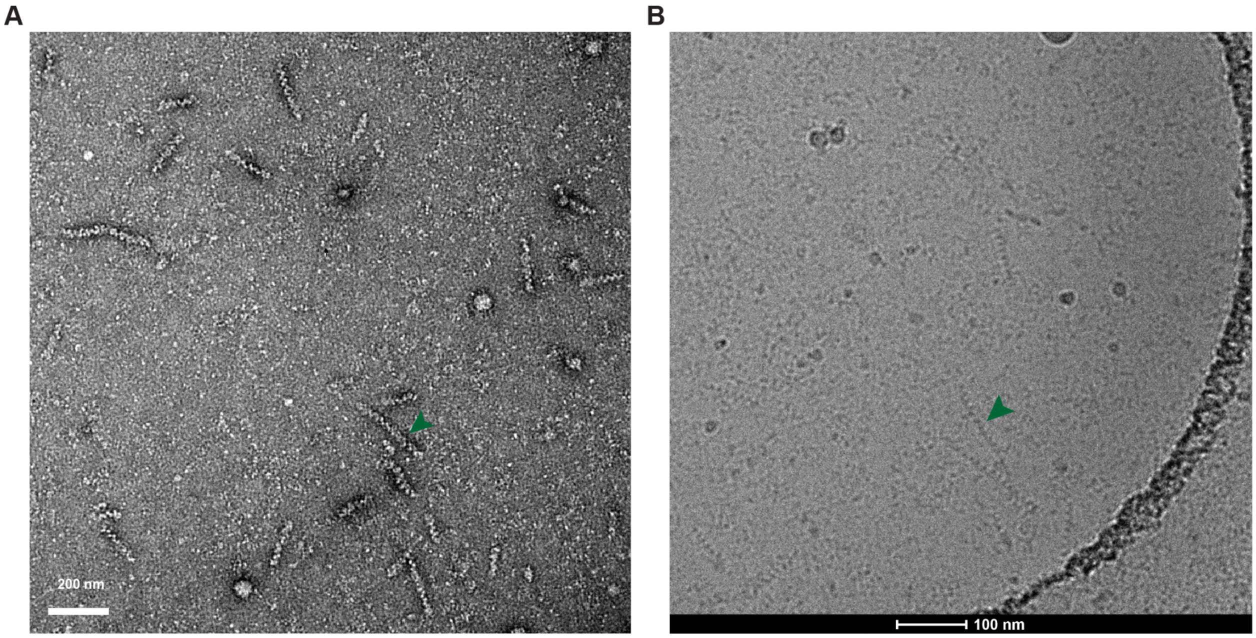


**Supplemental Figure 6.** Electron microscopy screening of enriched filament. Negative stain (A) and cryoEM (B) reveal an enrichment of filamentous assemblies and removal of larger vesicles and aggregates. Filaments are denoted by green arrows.

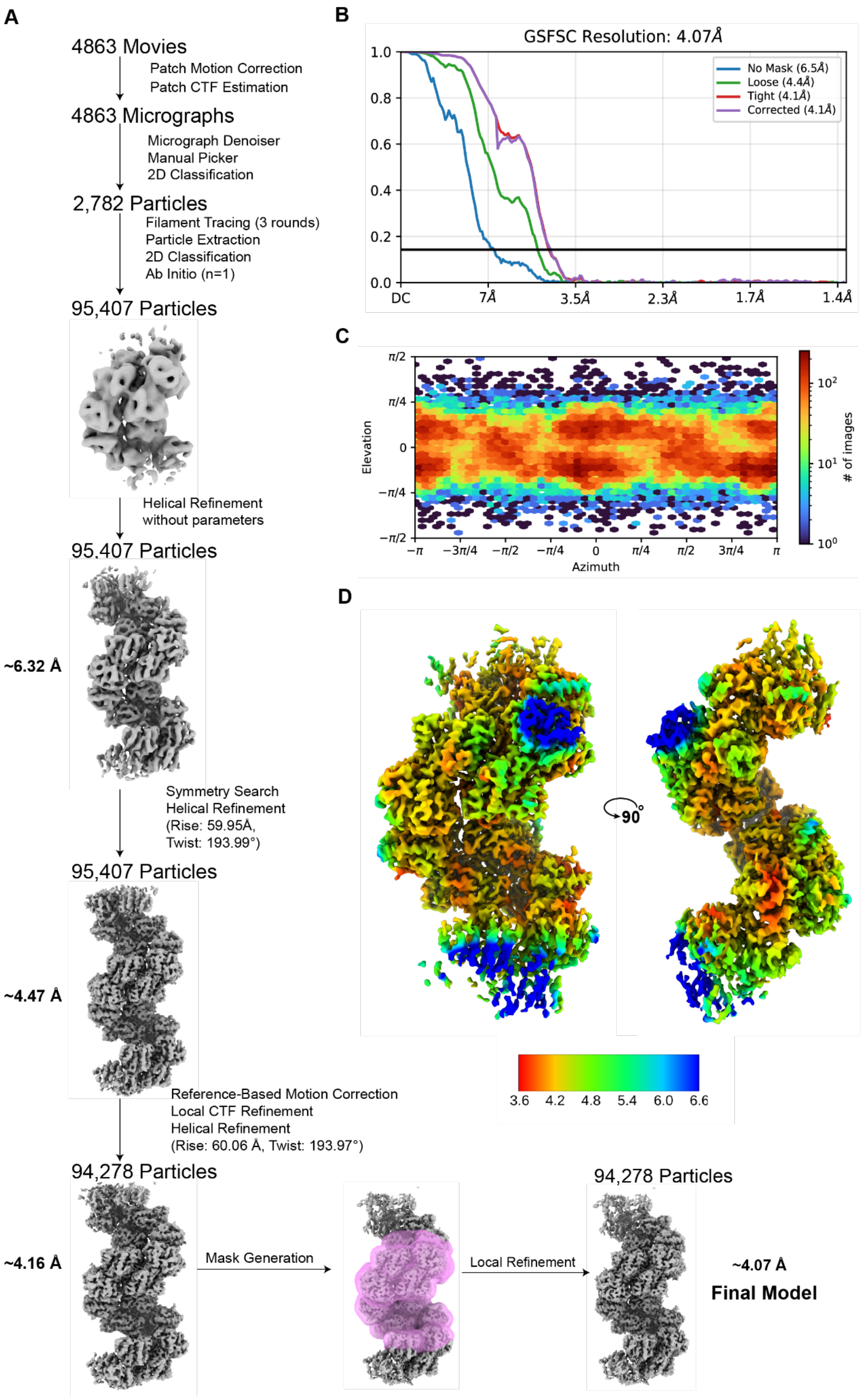


**Supplementary Figure 7.** Single-particle cryo-EM data processing pipeline for enriched filament fraction, resulting in a final helically refined map at ~4.07 Å resolution. Arrows summarize processes carried out in cryoSPARC v4, particle numbers and resolutions are displayed for notable density maps.

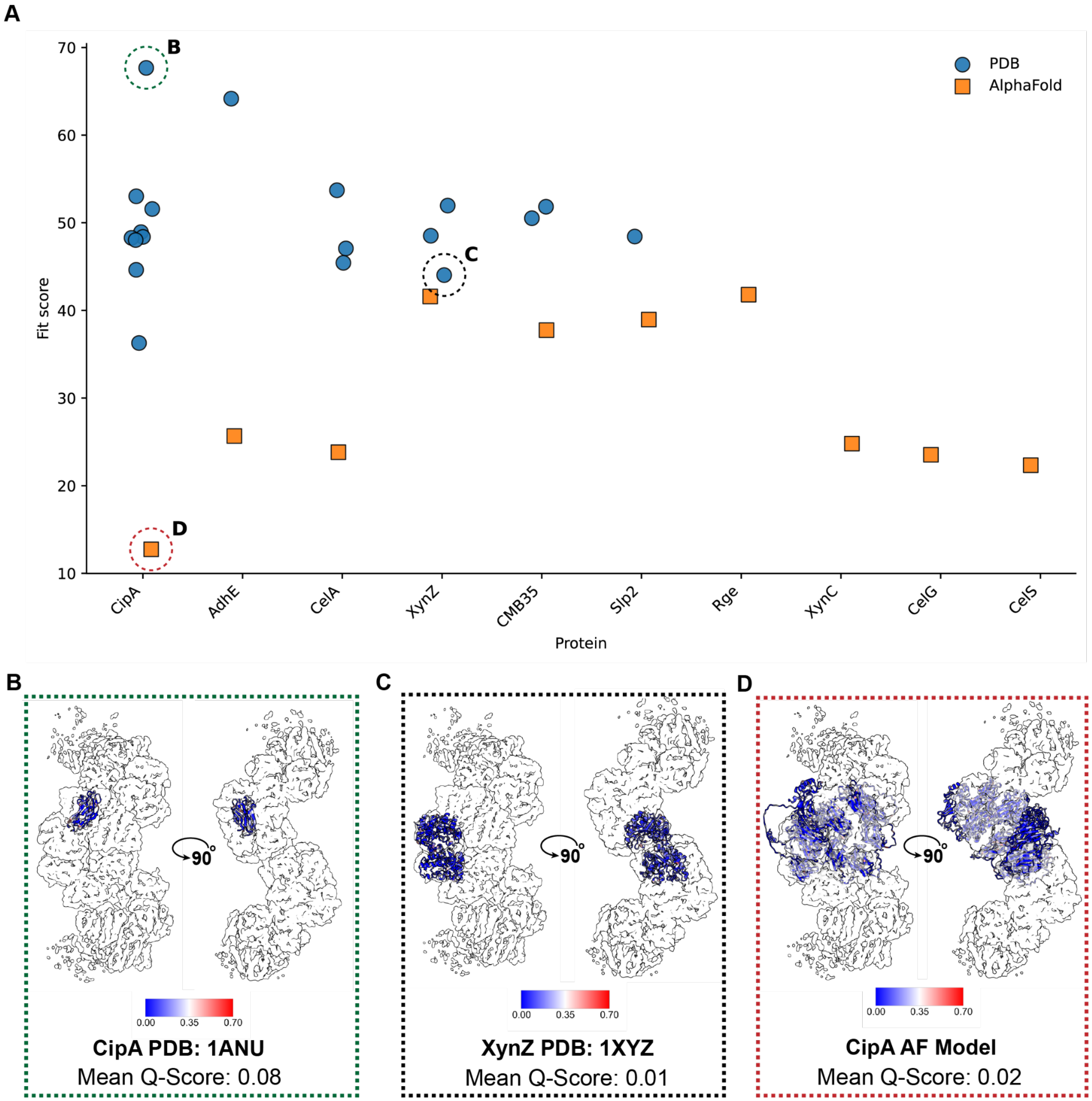


**Supplementary Figure 8.** Model fitting of proteomics-identified proteins. (A) All proteins were graphed by fit scores of either the PDB or AlphaFold-predicted models. The top fitting model by manual inspection - cellulosomal scaffold protein A (PDB: 1ANU) (B), medium fit model - endo 1,4 beta xylanase (PDB: 1XYZ) (C), worst model fit - cellulosomal scaffold protein A (AlphaFold2) (D).

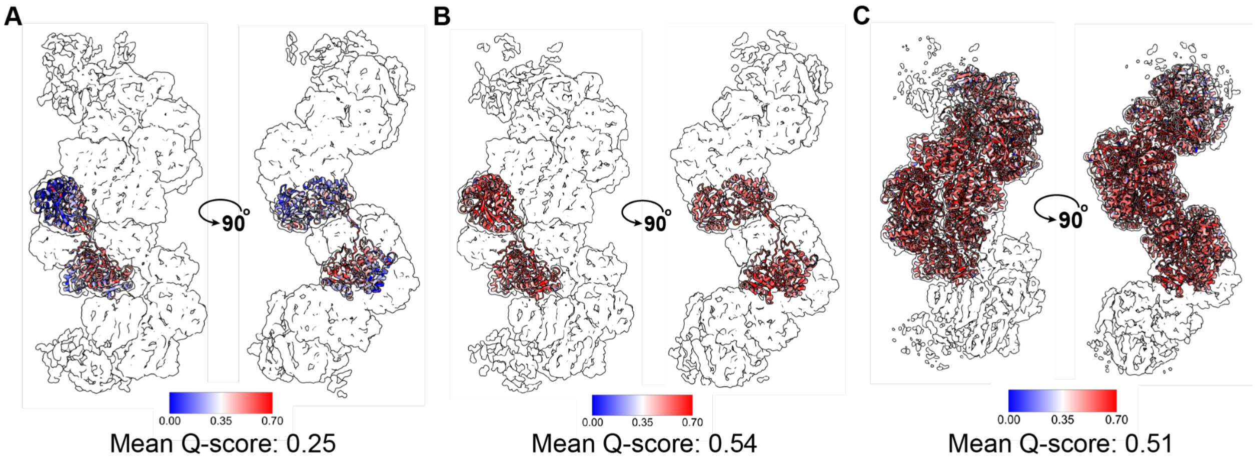


**Supplementary Figure 9.** Q-score analysis of AdhE. (A) Monomer of published *Ct* AdhE structure (PDB: 8UHW). Rosetta fast relax models of AdhE monomer (B), and full AdhE assembly (C).

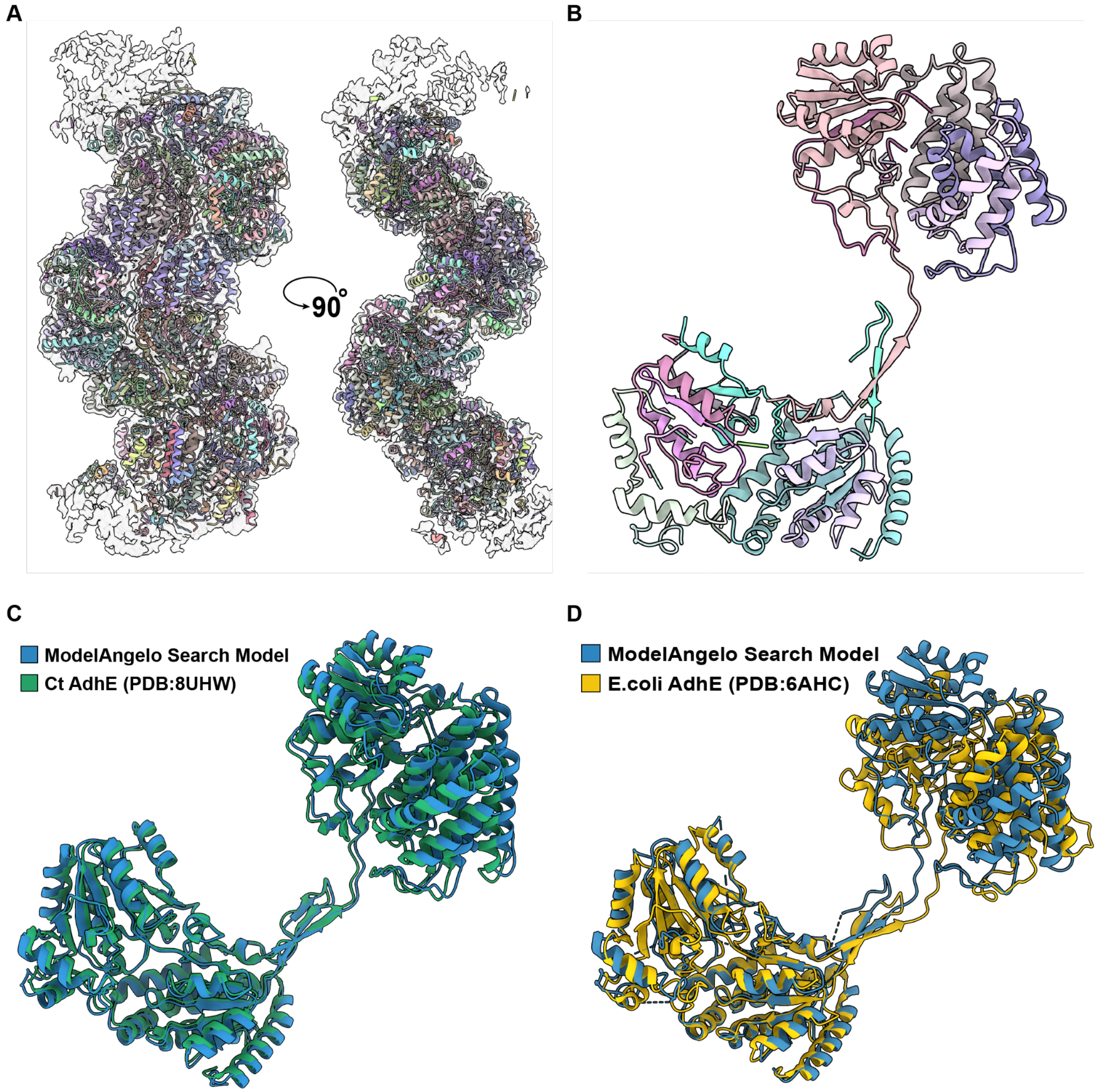


**Supplemental Figure 10.** Automated model building. (A) ModelAngelo fragments from automated model building in refined helical density. (B) Search monomer of ModelAngelo fragment output. (C) Top DALI structural search result and (D) top FoldSeek result, aligned onto ModelAngelo search model.

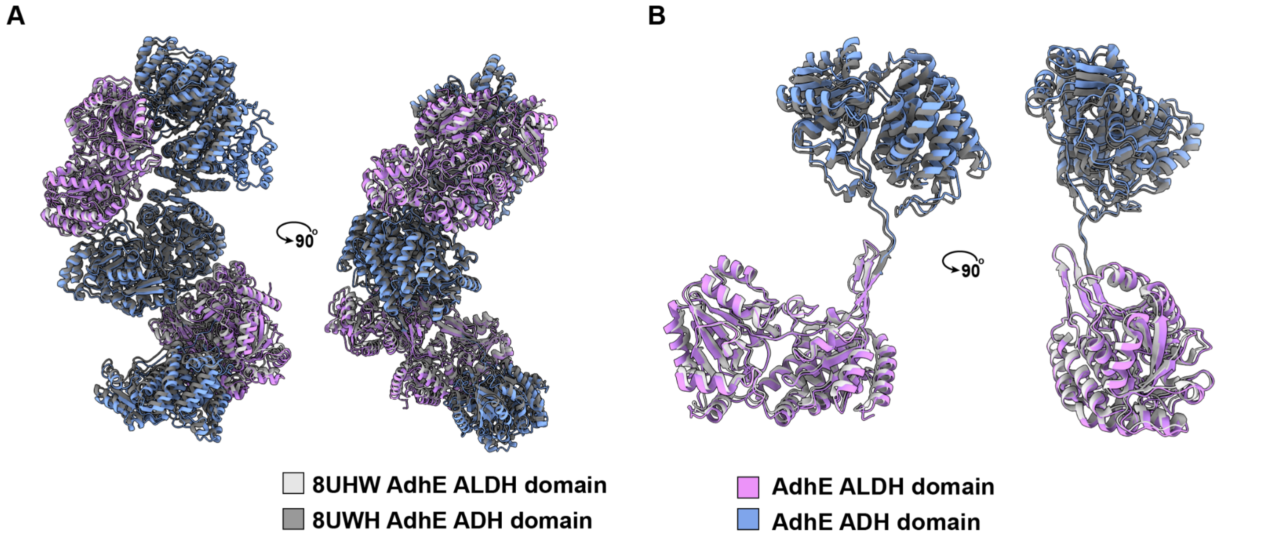


**Supplemental Figure 11.** Comparison of *C. thermocellum* AdhE structures. Overlay of entire AdhE model (A), and individual monomer (B) onto published *Ct* AdhE structure (PDB: 8UHW) used as initial model for structure refinement.

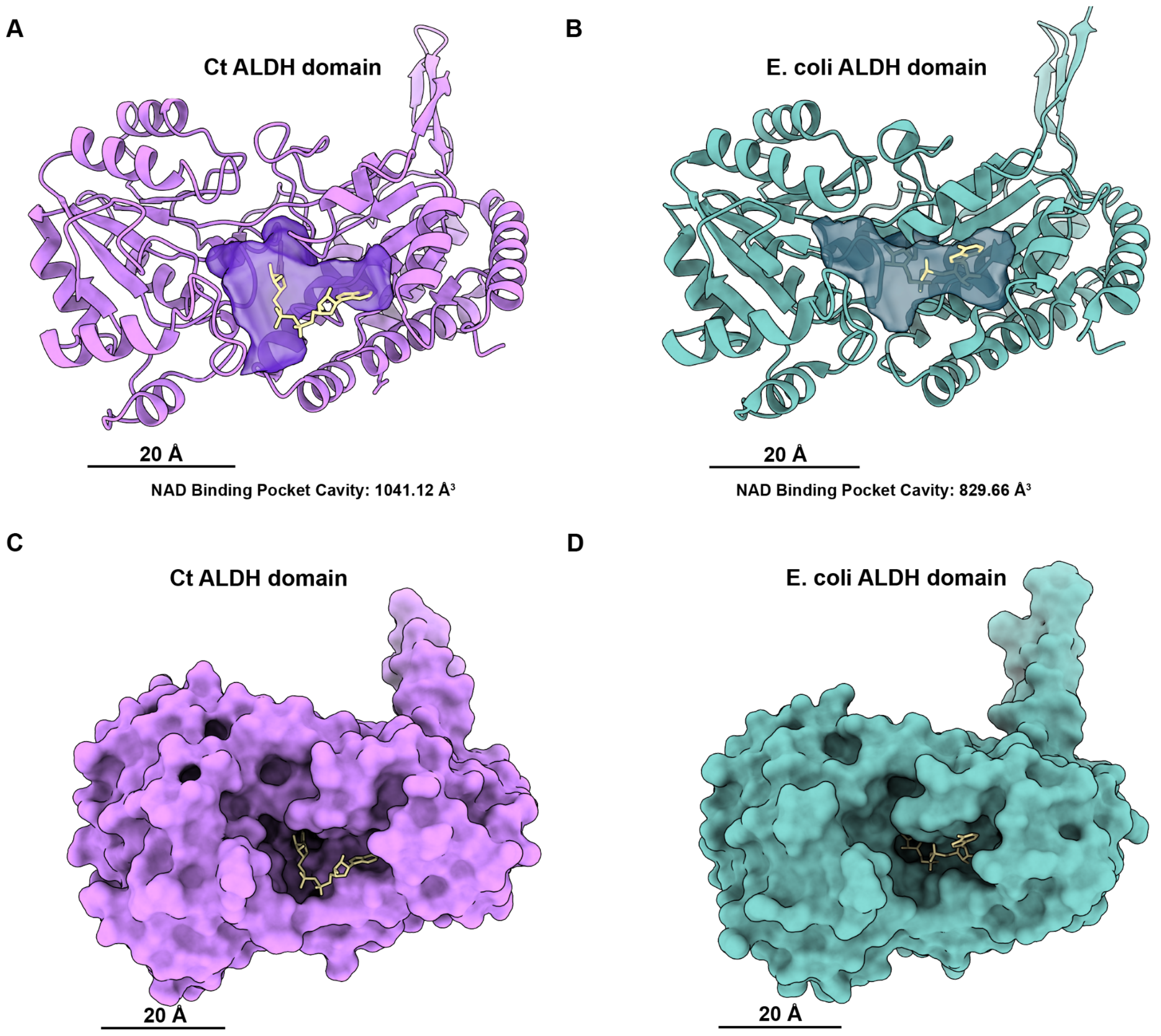


**Supplemental Figure 12.** Analysis of ALDH binding pocket. (A) *Ct* ALDH domain with bound $NAD^+$ depicting size of binding pocket. (B) Homologous ALDH domain from *E. coli* (PDB: 7BVP) with bound $NAD^+$ showing difference in cavity volume from *Ct* structure. Surface representations of each domain with bound $NAD^+$ illustrating depth of each protein's surface-accessible ligand cavity (C,D).

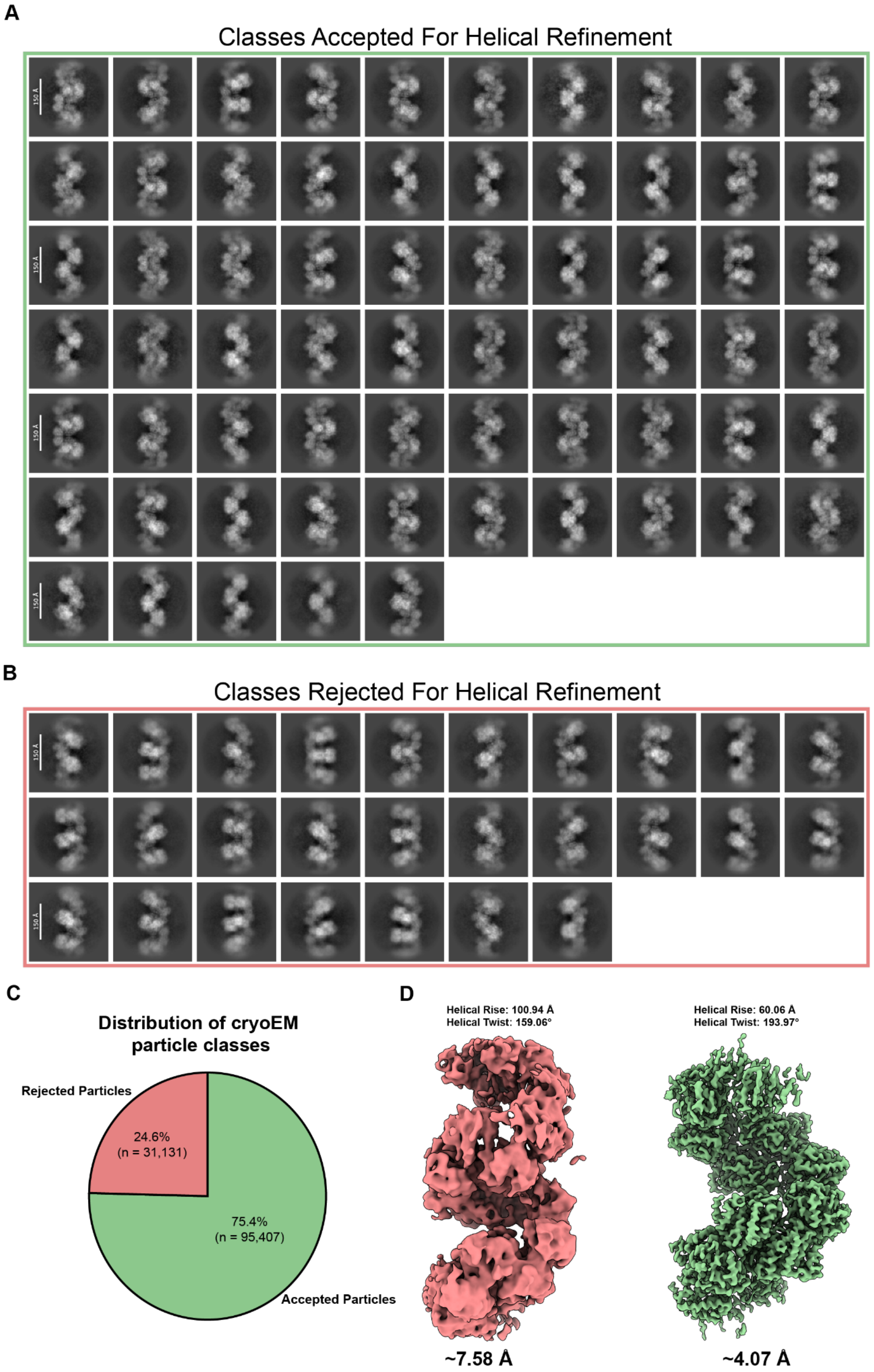


**Supplementary Figure 13**. Filament conformations observed in cryoEM data. (A) Views of extended conformation used in final filament structure determination. (B) Compact views were excluded from the processing pipeline. (C) Distribution of conformations observed in the full dataset broken up by classes used and classes rejected in final model processing. (D) Comparison between refined maps from rejected and accepted particle picks.

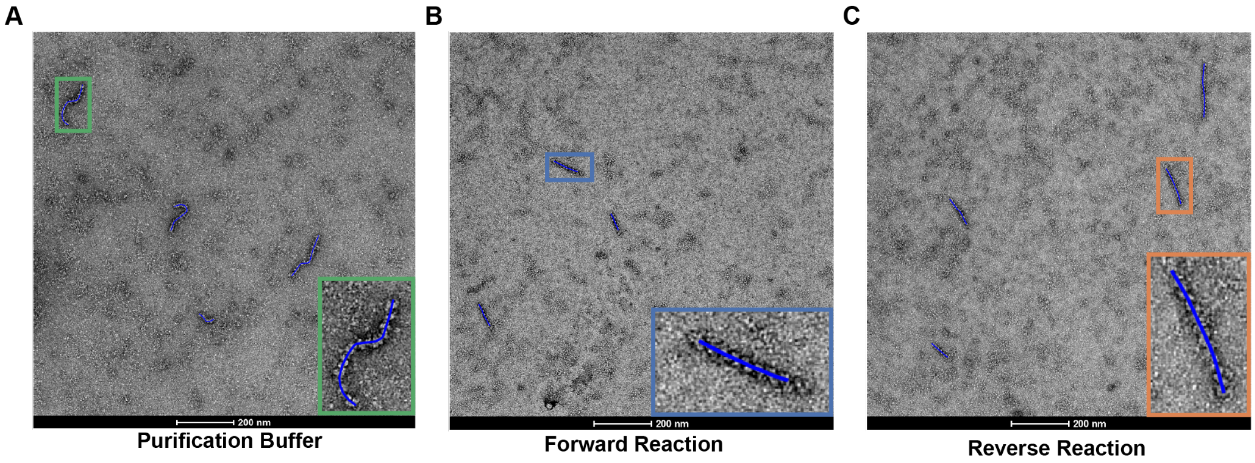


**Supplemental Figure 14.** FiberApp analysis of conformational changes. Example tracing for all three reaction conditions tests: the purification buffer (A), forward (B), and reverse reaction (C).

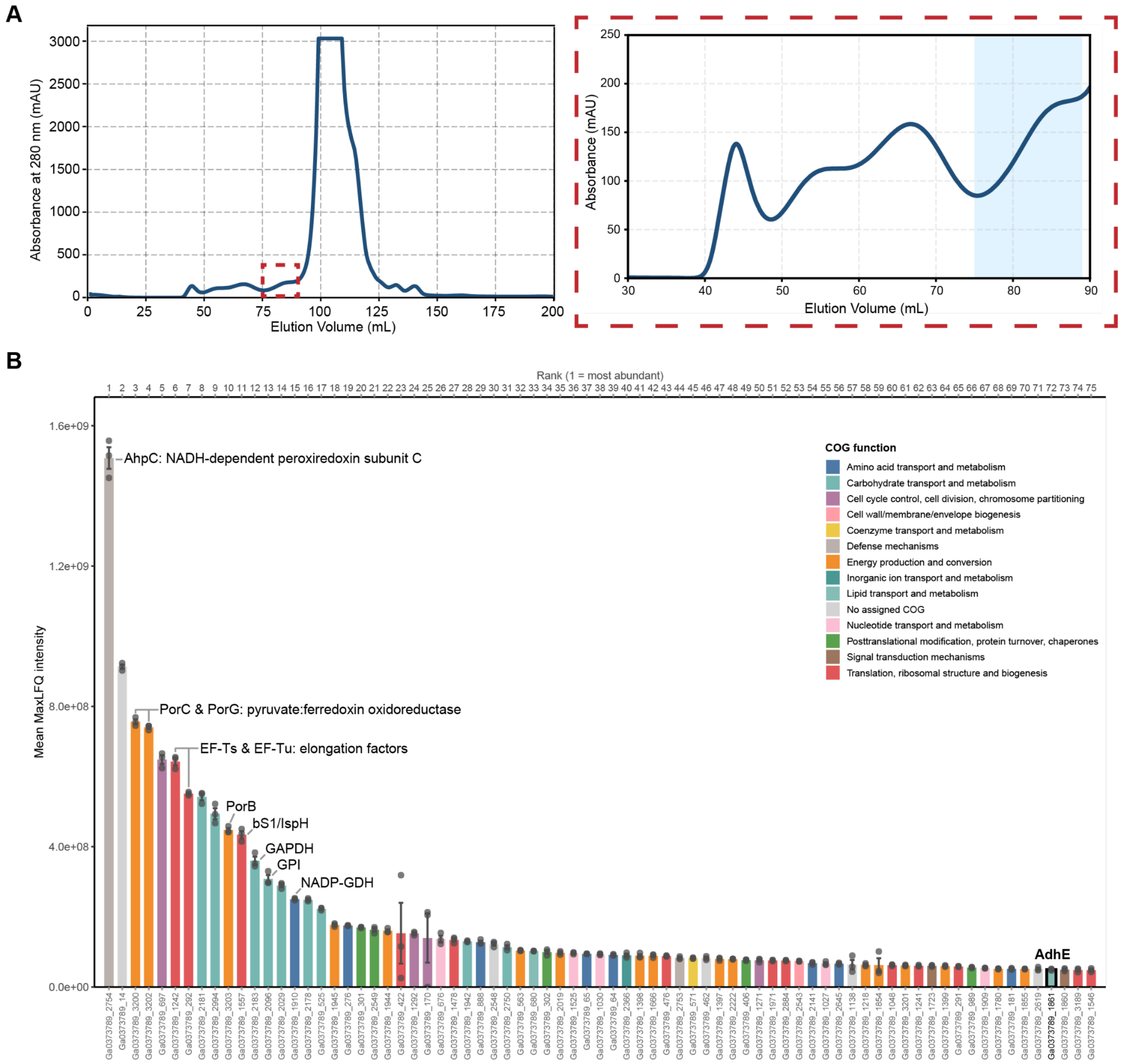


**Supplemental Figure 15.** Analysis of cytoplasmic lysis markers. Fractions corresponding to smaller molecular weight species from SEC fractionation (A), were analyzed by bottom-up proteomics and reveal presence of various cytoplasmic and nuclear proteins (B).
